# Attractor-based models for sequences and pattern generation in neural circuits

**DOI:** 10.1101/2025.03.07.642121

**Authors:** Juliana Londono Alvarez, Katie Morrison, Carina Curto

## Abstract

Neural circuits in the brain perform a variety of essential functions, including input classification, pattern completion, and the generation of rhythms and oscillations that support functions such as breathing and locomotion. There is also substantial evidence that the brain encodes memories and processes information via sequences of neural activity. Traditionally, rhythmic activity and pattern generation have been modeled using coupled oscillators, whereas input classification and pattern completion have been modeled using attractor neural networks. Here, we present a theoretical framework that demonstrates how attractor-based networks can also generate diverse rhythmic patterns, such as those of central pattern generator circuits (CPGs). Additionally, we propose a mechanism for transitioning between patterns. Specifically, we construct a network that can step through a sequence of five different quadruped gaits. It is composed of two dynamically distinct modules: a “counter” network, that can count the number of external inputs it receives via a sequence of fixed points; and a locomotion network, that encodes five different quadruped gaits as limit cycles. A sequence of locomotive gaits is obtained by connecting the counter network with the locomotion network. Specifically, we introduce a new architecture for layering networks that produces “fusion” attractors, binding pairs of attractors from individual layers. All of this is accomplished within a unified framework of attractor-based models using threshold-linear networks.

## 1 Introduction

Many neural circuits in the brain exhibit rhythmic or sequential activity, including cortical [1, 2, 3], hippocampal [4, 5, 6], and central pattern generator (CPG) circuits [7, 8]. At the same time, these brain regions also generate static patterns of persistent activity that support working memory, associative memory and pattern completion [9, 10, 11, 12]. Attractor neural networks (ANNs) have provided a useful paradigm for understanding *static* activity patterns. For example, ring and line attractors have been used for coding of head direction [13, 14] and eye position [15], and 2-dimensional bump attractor networks for place and grid cell coding [16, 17, 18, 19, 20, 21, 22]. This framework has also been used extensively to model memory encoding and retrieval [23, 24, 25, 26]. In all these cases, persistent activity is modeled with fixed point attractors – either as isolated discrete fixed points (e.g., for pattern completion) or as continuous families of fixed points (e.g., ring and line attractors). However, dynamic attractors such as limit cycles have remained under-explored. By incorporating a wider variety of attractors, can the ANN framework be further extended to develop models of rhythmic and sequential activity?

To date, rhythmic activity has traditionally been modeled using networks of coupled oscillators, assuming intrinsically oscillating or “pacemaker” neurons [27, 28, 29, 30, 31, 32, 33]. However, while there is evidence for pacemaker-type neurons in some CPGs, it has been noted that for many brain rhythms, including those underlying locomotion, network interactions mediated by inhibition play a more prominent role [34]. Analogously, it has been argued that cortical oscillations arise from an interplay between inhibition and network connectivity [3]. Another limitation of coupled oscillator models, particularly for locomotion, is that they often require fine-tuned parameters, and changes of parameters are necessary to transition between different oscillatory patterns (such as different gaits).

Here, we investigate the use of dynamic attractors for rhythmic and sequential activity, thus bringing these phenomena into a unified attractor neural network framework. Dynamic attractors include limit cycles, quasiperiodic and chaotic attractors, but here we primarily focus on limit cycles. These are natural candidates to model oscillatory activity that is robust to perturbations. Our mathematical setting is threshold-linear networks (TLNs), as these have been shown to support both static and dynamic attractors [35, 36, 37]. TLNs have been widely used in computational neuroscience for modeling via fixed point attractors [26, 38, 39, 40, 41], and here we extend the framework to modeling rhythmic activity with dynamic attractors.

TLNs model the firing rates, *x*_*i*_(*t*), of *n* recurrently connected neurons that evolve in time according to the equations:

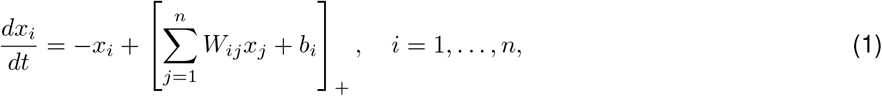

where [·]_+_ = max {0,·} is the threshold nonlinearity. A given TLN is specified by the choice of a connection strength matrix *W* and a vector of external inputs *b* ∈ ℝ^*n*^. Importantly, all the fixed point attractor phenomena (including ring, line, and 2-dimensional bump attractors) are reproducible with inhibition-dominated TLNs [18, 42, 13, 14, 20], as is Hopfield-style associative memory encoding and retrieval [25, 42, 43, 44, 45, 46]. In each of these cases, there are explicit connectivity matrices *W* that have been found to yield the desired stable fixed point attractors. These originated as theoretical ansatzes, without knowledge of the true connectivity between neurons, but recent advances in connectomics have shown that some of these connectivity structures do in fact appear in the brain [47]. Can we similarly understand what types of connectivity structures give rise to dynamic attractors?

Until recently, relatively little was understood about how to identify or engineer TLNs with a prescribed set of dynamic attractors, and thus modeling rhythmic and sequential activity in this setting remained limited. But this has begun to change with the introduction of a special subfamily of TLNs, known as *combinatorial* threshold-linear networks (CTLNs), whose connectivity matrices are dictated by a simple directed graph [48]. Several theoretical results on these networks show how features of the underlying connectivity graph shape the network’s dynamics [35, 36, 37, 45, 49, 50, 51]. This has enabled the design and identification of CTLN motifs that exhibit various patterns of rhythmic activity (dynamic attractors). Remarkably, these dynamic attractors can coexist alongside the more traditional stable fixed point attractors.

In this work, we illustrate these ideas and provide a proof of concept that the ANN framework can be extended to models of rhythmic activity, including multiple coexisting dynamic attractors in the same network. Specifically, we present a CPG network that can robustly encode five prescribed quadruped gaits. Each gait corresponds to a distinct limit cycle driving the same four “limb” neurons in a different pattern. All five limit cycles are accessible

– without changing parameters – simply by choosing different initial conditions or providing brief control pulses to induce gait transitions. This contrasts with most existing locomotion models, where network parameters such as synaptic weights must be adjusted to switch between gaits.

We then asked if a sequence of gaits could be stored in a network, such that identical input pulses could trigger transitions in a prescribed order. To engineer such a model, we first devised a discrete counter/neural integrator network that encodes the total number of received pulses via the position of a bump of activity within a chain of stable fixed point attractors. This network exploits the fact that CTLNs can easily store multiple stable fixed points [45, 46], together with a simple hysteresis mechanism. Next, we combined the counter and quadruped gaits networks so that identical pulses to the counter network resulted in a prescribed set of transitions between quadruped gaits. By changing the wiring between the counter and an intermediate “relay” layer, different sequences of gaits can be stored while leaving the gait subnetwork intact.

The attractors of this combined network are neither the fixed points of the counter nor the original limit cycles for the gaits. Instead, they are *fusion attractors*, binding together gaits and sequence position in all possible combinations. The same gait can thus be flexibly reused multiple times in a single prescribed sequence, since it can be fused with any of the fixed points representing sequence position in the counter network. Notably, the timing of the transitions between gaits is not determined by the network architecture, but is externally driven by the timing of input pulses. Thus, sequence order, sequence timing, and the temporal pattern of activity within each gait are all dissociated and controlled by different aspects of the network.

Our work provides not only a proof of concept that rhythms and oscillations can be modeled within an ANN framework, but also suggests concrete mechanisms for building modular networks that combine and reuse both static and dynamic attractors. In particular, the simple architecture of the gait-counter network, and the novel binding mechanism of fusion attractors, has implications for understanding how the brain (or a robot) may organize complex behaviors such as a choreographed sequence of dance moves.

## 2 Results

### 2.1 TLNs as a unified framework for static and dynamic attractors

We begin by outlining the theoretical framework underlying this work. As mentioned before, we use CTLNs: graph-based, inhibition-dominated TLNs in which the graph reflects connections between excitatory neurons in the presence of global inhibition. This framework is both mathematically tractable and dynamically expressive. Indeed, there is a large body of theoretical work that directly relate features of the graph to features of the network dynamics [35, 36, 37, 45, 49, 50, 51], making it possible to analyze and design networks that exhibit a wide range of dynamic behaviors [37].

#### TLNs built from graphs

Combinatorial threshold-linear networks (CTLNs) are a special family of TLNs that are parametrized by a triple (*G, ε, δ*), where *G* is a simple directed graph and *ε, δ >* 0 are real-valued parameters. The connectivity matrix *W* = *W* (*G, ε, δ*) is defined as follows:

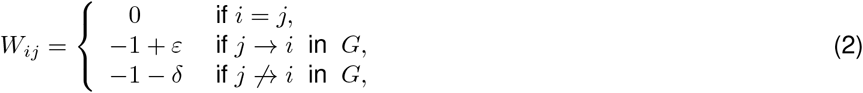

and the external input *b* is typically taken to be uniform across neurons and constant in time: *b*_*i*_(*t*) = *θ* for all *i* = 1, …, *n*. When 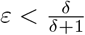, we say the parameters are in the *legal range*.^1^ Throughout this work, we will assume parameters fall within the legal range unless explicitly stated otherwise; in particular, all simulations in the main text were done with *ε* = 0.25, *δ* = 0.5. We need this assumption on the legal range for most of the theoretical results used here. Additionally, we assume that all CTLNs considered are *nondegenerate*.^2^

Note that the upper bound on *ε* implies *ε <* 1, and so the *W* matrix is always effectively inhibitory. Heuristically, we think of CTLNs as modeling a network of excitatory neurons within a sea of global inhibition, so that the *W*_*ij*_ values represent the sum of excitatory and inhibitory weights. Recently, this heuristic was made precise: it was shown that every CTLN is equivalent to an excitatory-inhibitory (E-I) network with the same connectivity graph and a single inhibitory node [52, 53]. When the timescale of inhibition is sufficiently fast, the E-I network matches the dynamics of the CTLN and the attractors appear to be identical (see Supplementary Figure S1). CTLNs have also recently been shown to be a useful mean-field approximation for inhibition-stabilized spiking networks with clustered excitatory connectivity [52].

Figure 1A-C illustrates the variety of static and dynamic attractors that TLNs can exhibit. Panel A shows the graph of a CTLN together with a solution of the dynamics that converges to a stable fixed point that is a global attractor of this network. Panel B shows a CTLN that produces rhythmic activity via a limit cycle. Finally, panel C shows an example CTLN that produces a fusion attractor in which the activity at a fixed point is bound together with that of a simple limit cycle.

**Figure 1:**
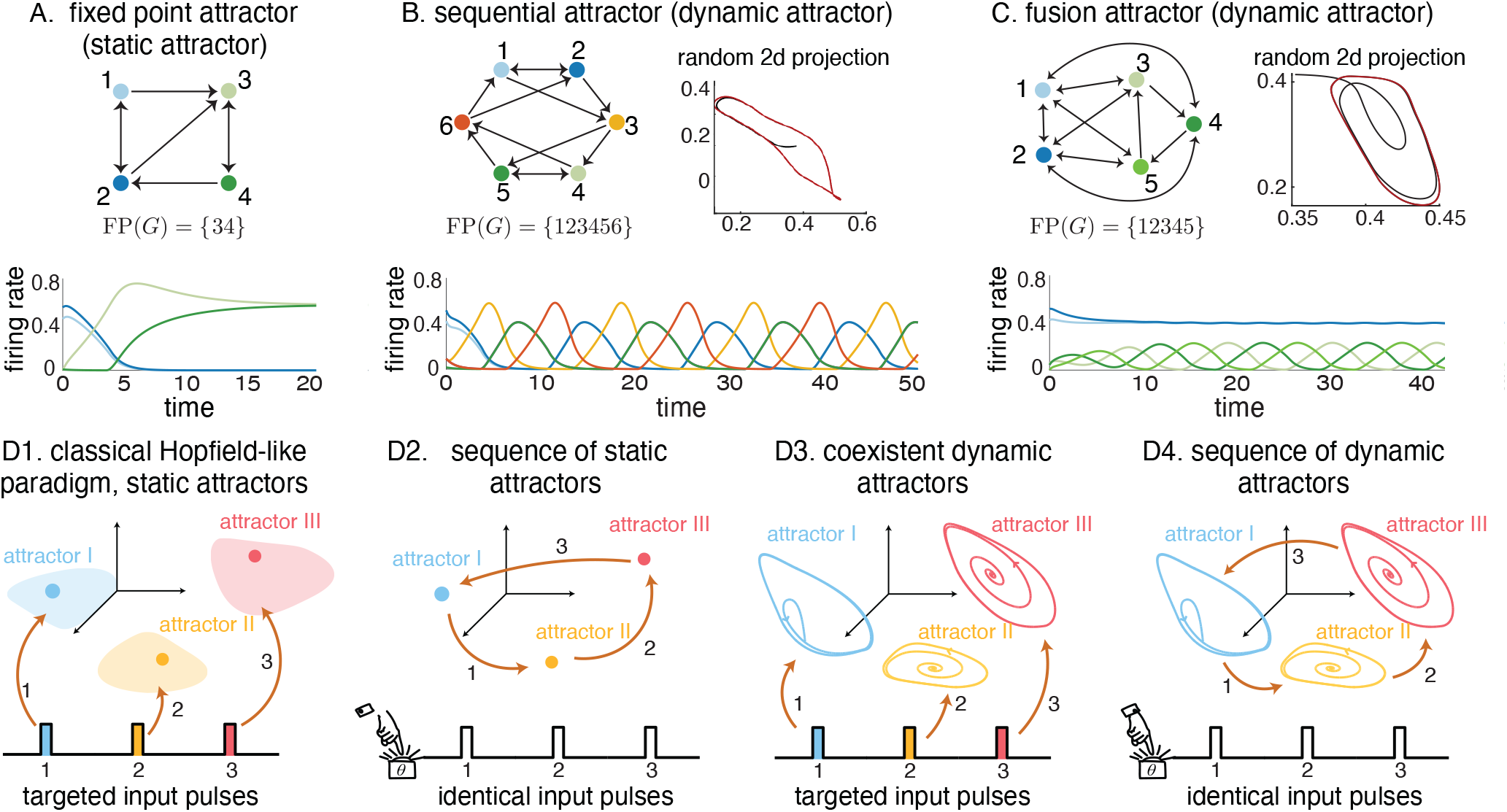
Attractor types and sequences of attractors. (A-C) Top: Graphs defining CTLNs. Bottom: Firing rate curves *x*_*i*_(*t*) for particular solutions of the corresponding CTLN above, matching the color of the nodes. (A) Fixed point attractor corresponding to a clique in the graph of a CTLN. (B) Sequential attractor (limit cycle) of a CTLN whose graph is a *cyclic union*. Neurons take turns firing in sequence, with neurons 1 and 2 firing synchronously, as well as neurons 4 and 5. (C) Fusion attractor arising from a *clique union*. The attractor fuses a fixed point and a limit cycle. (B-C, top right) The trajectory is shown in a random 2-dimensional projection, with the last three quarters in red to highlight that the attractor is a perfectly periodic limit cycle. (D1) Classical Hopfield-like paradigm: multiple stable fixed points (static attractors) are encoded in the same network, each accessible via attractor-specific (colored) input pulses that push the network state into the corresponding basin of attraction (colored blobs). (D2) Multiple stable fixed points are encoded in the same network, each accessible in a predetermined sequence via identical pulses. (D3) Multiple *dynamic* attractors are encoded in the same network, each accessible via attractor-specific (colored) input pulses. (D4) Internally encoded sequence of dynamic attractors, with each step of the sequence triggered by identical inputs.

#### Prior results relating connectivity to dynamics

There is a large body of theoretical results connecting graph structure to CTLN dynamics [35, 36, 37, 45, 49, 50, 51, 53]. Most of these results give constraints on the collection of stable and unstable fixed points of the network in terms of features of the graph. In addition, there is a body of computational work showing that the fixed point structure is predictive not only of static attractors (stable fixed points), but also of the dynamic attractors [49]. In particular, each dynamic attractor typically has an *unstable* fixed point associated to it in the following sense: (1) initial conditions near this fixed point yield trajectories that converge to the attractor, and (2) the high-firing neurons of the attractor correspond to the active neurons at the fixed point. Thus we track both the stable and unstable fixed points of a CTLN through the combinatorial object FP(*G*) defined below.

The *support* of a fixed point *x*^*\**^ is the subset of its active neurons: 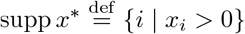. For a nondegenerate TLN, there can be at most one fixed point per support [45]. Thus, we can label all the fixed points of a network by their support, *σ* = supp *x*^*\**^ *⊆* [*n*], where 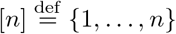. For a CTLN, we denote this collection of supports by

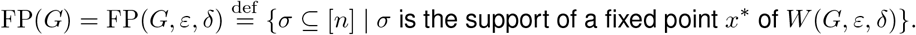

All *graph rules* are independent of the choice of parameters *ε* and *δ*, as long as these fall within the legal range, and thus we suppress these parameters from the notation FP(*G*) for convenience. From FP(*G*) it is straightforward to recover the precise values of the fixed points: given a support *σ* ∈ FP(*G*), the corresponding fixed point *x*^*\**^ satisfies 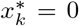 for all *k* ∉ *σ* and 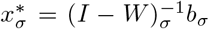, where the subscript *σ* denotes restriction of the vector or matrix to the indices in *σ*.

Below we review the graph rules (theorems) needed here, beginning with the graph structures that give rise to stable fixed points. Recall that for a subset of nodes *σ ⊆* [*n*], the induced subgraph *G*| _*σ*_ is a *clique* if it is all-to-all bidirectionally connected.

#### Rule 1 (cliques [45, 35])

If *G*|_*σ*_ is a clique, then *σ* ∈ FP(*G*) if and only if there is no node *k* of *G, k* ∉ *σ*, such that *i* → *k* for all *i* ∈ *σ*. In other words, *σ* ∈ FP(*G*) if and only if *G*|_*σ*_ is a *target-free* clique. If *σ* ∈ FP(*G*), the corresponding fixed point is stable.

For example, in Figure 1A, both *σ* = {1, 2} and *τ* = {3, 4} are cliques, but *σ* has a target (node 3), so it does not support a fixed point in the full network. Note that within its restricted subnetwork, *G*| _*σ*_, the clique *does* support a fixed point, but that fixed point *dies* within the larger network with the addition of node 3. In contrast, *τ* is target-free so it does support a fixed point, and this fixed point is stable. We say that fixed point for *τ survives* in the larger network.

It has been conjectured that target-free cliques are the only graphs that yield stable fixed points in CTLNs [45, 46]. If the conjecture holds, then any graph that has no target-free cliques would necessarily yield a CTLN where all the fixed points are unstable, and thus only dynamic attractors can emerge. We see this in the CTLNs in panels B and C of Figure 1: each network has a single fixed point, which is unstable. The networks also each have a single global attractor, which is dynamic.

Rule 1, which characterizes stable fixed points via target-free cliques, is a special case of the uniform in-degree rule. This more general result applies to any subgraph *G*|_*σ*_ where each node receives exactly *d* incoming edges from within *G*|_*σ*_. In that case we say that *G*|_*σ*_ has uniform in-degree *d*. These subgraphs always support a fixed point in *G*|_*σ*_, and the theorem below gives conditions under which the fixed point survives in the full network:

**Theorem 1** (uniform in-degree, [45, 35]). *Suppose G*|_*σ*_ *has uniform in-degree d, then σ* ∈ FP(*G*|_*σ*_). *For each k* ∉ *σ, let* 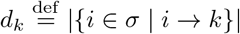 *be the number of edges k receives from σ. Then*

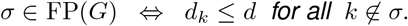

*Furthermore, if* |*σ*| *>* 1 *and d <* |*σ*| */*2, *then the fixed point is unstable. If d* = |*σ*| − 1, *i*.*e. if σ is a clique, then the fixed point is stable*.

For instance, the graph in Figure 1B has uniform in-degree 2, and thus the network has a full support fixed point. On the other hand, the subgraph *G*| _{3,4,5}_ in panel C has uniform in-degree 1, but since node 1 receives edges from that subgraph, {3, 4, 5} does not survive to yield a fixed point of the full network. Similarly, for the clique supported on {1, 2}, the fixed point does not survive in the full network.

The graphs in Figure 1B and C have additional structure that gives further insight into the attractors they generate. The graph in B is known as a *cyclic union* in which component subgraphs are connected together in a cyclic fashion. These networks typically produce sequential attractors, and the structure of FP(*G*) has been fully characterized [45, 51]; we review this in Section 2.3 where we use this type of network to produce quadruped gaits. The graph in panel C is known as a *clique union* as its component subgraphs *G*| _{1,2}_ and *G*| _{3,4,5}_ are all-to-all connected to each other via cliques. This is one construction known to produce fusion attractors and whose FP(*G*) has also been fully characterized [45]. In Section 2.4, we develop another graph construction via a relay layer that also yields fusion attractors, and we characterize its collection of fixed points and attractors.

As noted before, there is significant computational evidence that dynamic attractors of CTLNs are associated with particular *unstable* fixed points of the network [49, 35]; specifically, the unstable fixed points come from minimal subnetworks, known as *core motifs*. We say that *G*| _*σ*_ is a *core motif* if it has a full support fixed point and no proper subsets yield surviving fixed points, i.e. FP(*G*| _*σ*_) = {*σ*}. Given the strong connection to attractors, we are particularly interested in the subset of FP(*G*) consisting of core motifs:

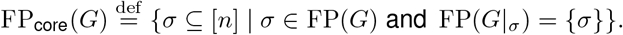

Note that a network can have multiple core motifs, yielding both stable and unstable fixed points, which can result in the coexistence of multiple static and dynamic attractors [49, 36, 37].

In general, switching between attractors in such systems is a nontrivial problem, as nonlinear dynamical systems often have complicated basins of attraction [54, 55]. However, for CTLNs, this challenge can be easily addressed through the external input *b* parameter. Although the theoretical results above apply to networks with uniform, constant external input (*b*_*i*_(*t*) = *θ >* 0 for all *i*), in the remainder of this work, we extend this framework to allow piecewise constant inputs in the form of brief pulses, in which *b*_*i*_(*t*) is temporarily increased for some or all neurons during a short time interval. These step inputs effectively define a sequence of piecewise CTLNs, and can be used to perturb the system in order to switch between attractors.

#### Sequences of attractors: external vs internal encoding

Figure 1D shows schematics of different types of sequences of static and dynamic attractors that are accessed via brief input pulses. In panel D1, multiple stable fixed point attractors coexist, with transitions between them requiring attractor-specific inputs (as in Hopfield networks). That is, each external pulse must target the neurons of the next attractor in the sequence, making the sequence *externally* encoded. In contrast, panel D2 shows an *internally* encoded sequence of fixed point attractors, where identical input pulses sent to the full network trigger transitions, and the sequence order is stored in the network connectivity rather than the input. This kind of mechanism could be useful for modeling highly stereotyped sequences, and underlies the function of the counter network, which we present next in Section 2.2. Panels D3 and D4 illustrate the dynamic attractor analogues of these two cases. In panel D3, multiple dynamic attractors (e.g., limit cycles) coexist in state space, with transitions between them requiring attractor-specific inputs, as will be the case for the quadruped gaits network (Section 2.3). In panel D4, the sequence of dynamic attractors is internally encoded, and identical input pulses trigger transitions through a predefined order encoded by the network connectivity. The combined gaits-counter network is an example of this (Section 2.4).

Because static attractors are typically easier to engineer and analyze than dynamic attractors, we begin with the simplest case: a sequence of fixed point attractors (Fig. 1D2).

### 2.2 TLN counter

In this section we model a discrete counter as a sequence of stable fixed points, each accessible via identical inputs, as shown in Figure 1D2. Neural integration is an important correlate of processes such as oculomotor and head direction control, short term memory keeping, decision making, and estimation of time intervals [56, 57, 58, 59]. Classic neural integration models are known to be fine-tuned, requiring exact values for the parameters in order to achieve perfect integration. In contrast, some more robust models tend to be rather insensitive to weaker inputs, requiring strong inputs to switch between adjacent states [56, 58].

By Theorem 1, stable fixed points can be engineered via target-free cliques. We thus construct our network by chaining together several target-free cliques, as shown in Figure 2A. We have wrapped the chain around to ensure activity will not stall in the last clique of the chain. By Theorem 1 these 2-cliques will yield surviving fixed point supports as long as they do not have more than one outgoing edge to any other node in the graph. For instance, if we were to add the edge 1 → 4, then nodes {1, 2}, associated to the first clique, would no longer support a fixed point of the network, as node 4 would now be a target. We have thus built a network whose FP(*G*), by design, should contain all the cliques ({1, 2}, {3, 4}, {5, 6}, {7, 8}, {9, 10}, {11, 12}) as core fixed point supports. Note that each cycle ({1, 3, 5, 7, 9, 11}, {2, 4, 6, 8, 10, 12}) also supports a fixed point of the network by Theorem 1, since no node receives more than one edge from either cycle. Computationally, we confirm these inclusions and found, overall that | FP(*G*)| = 141 and

**Figure 2:**
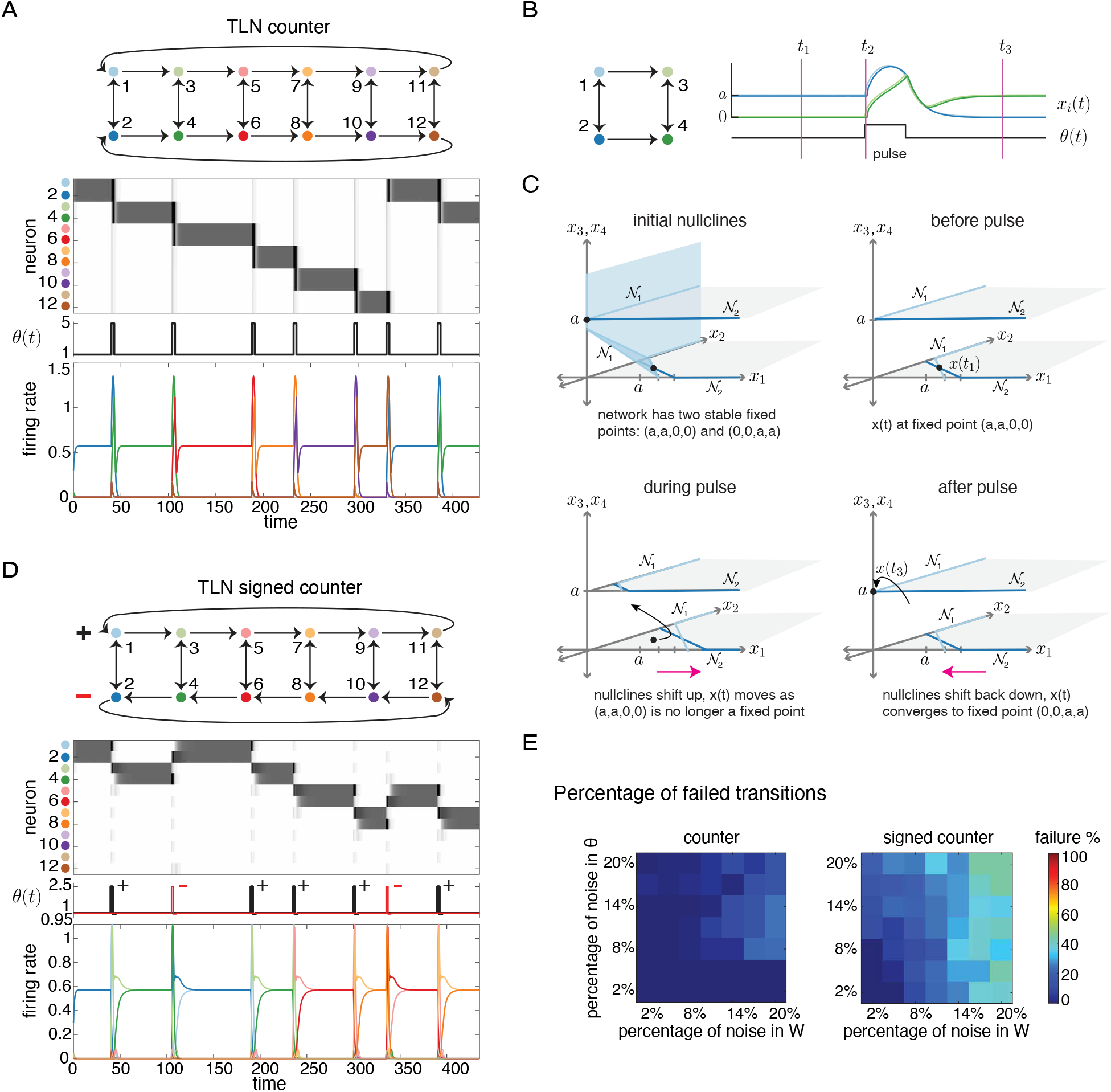
TLN counters. (A) Unidirectional chain of 2-cliques. Pulses are sent to all neurons in the network, sliding activity to the next clique. Pulse duration is 2 time units. (B) A small subnetwork consisting of only two 2-cliques. The state is initially in a stable fixed point supported on the 1 ↔ 2 clique (blue) with *x*(*t*) = (*a, a*, 0, 0) with 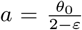. After the pulse, the network settles into the second stable fixed point, corresponding to the 3 ↔ 4 clique (green), with *x*(*t*) = (0, 0, *a, a*). (C) Illustration of the hysteresis mechanism underlying the fixed point transitions. (Top left) Nullclines for the first pair of nodes. The 𝒩_1_ nullcline is the set of states where 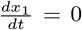. It consists of two hyperplane segments, one for *x*_1_ = 0 and one for *x*_1_ *>* 0 (shaded in blue). The 𝒩_2_ hyperplane has the same kind of structure but only the intersections with the horizontal slices *x*3, *x*4 = 0 and *x*3, *x*4 = *a* are shown for ease of visualization. The system has two stable fixed points, one on each of these slices (black dots). (Top right) Before the pulse the network activity is at the stable fixed point (*a, a*, 0, 0) supported on the 1 ↔ 2 clique. (Bottom left) A *θ*-pulse shifts the nullclines in state space, causing the trajectory (black) to move. (Bottom right) When the pulse ends, the network state *x*(*t*) is in a new basin of attraction and converges to the stable fixed point (0, 0, *a, a*). (D) A TLN signed counter is constructed by flipping the direction on the bottom cycle of even numbered neurons. Black pulses shift activity right (odd neurons), red pulses shift left (even neurons). Pulse duration is 2 time units, with 3 time units of refractory period during which *θ* = 0.95. (E) Failed transition percentage per pulse trial as a function of noise in *W* and *θ*. Heatmap shows failed transitions, with axes representing noise percentages in *θ* and *W*. For details on how it was computed see Supplementary Materials section A.2.

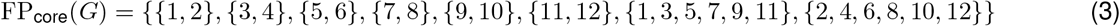

This indicates that we will have a stable fixed point attractor per clique, and two unstable fixed points giving rise to dynamic attractors. To test this prediction, we simulated the network using the parameters *ε* = 0.25, *δ* = 0.5, a baseline *θ* = 1, and an input pulse *θ* = 5 of 3 time units duration. However, since Theorem 1 does not depend on the values of these parameters, any TLN with dynamics prescribed by equation (1), whose connectivity matrix *W* is built from the graph of Figure 2A according to equation (2), will have the same fixed point structure (eq. (3)). This is true for any values of *ε* and *δ*, provided they are within the legal range.

The FP_core_(*G*) predictions about the dynamics are partially confirmed by the rate curves of Figure 2A: activity remains at the stable fixed point that the network was initialized to, until a uniform *θ*-pulse is sent to all neurons in the network. When the network receives the pulse, activity slides down to the next stable state in the chain. Activity is maintained in this state indefinitely until future pulses are provided to the system. Thus, the network is effectively counting the number of input pulses it has received via the position of the attractor in the linear chain of attractor states. The count is maintained via well-separated *discrete* states, providing a straightforward readout mechanism of the encoded count.

On the other hand, the supports {1, 3, 5, 7, 9, 11}, {2, 4, 6, 8, 10, 12} of equation (3) (corresponding to the top and bottom cycles) predicted the existence of two dynamic attractors, which were not found computationally.

Note that the value of the current fixed point depends on the value of the previous fixed point, so the network “remembers” where it came from (this is called *hysteresis*). How does this phenomenon arise from the *θ* pulse? In Figure 2B we present a short version of the counter, along with a detailed view of activity around the pulse. Hysteresis occurs because fixed points lie at intersections of *nullclines* 𝒩_*i*_ (subsets of the state space where 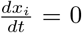, and the arrival of a pulse causes the nullclines to shift. The *i*-th nullcline of a TLN is the union of a pair of hyperplane segments given by

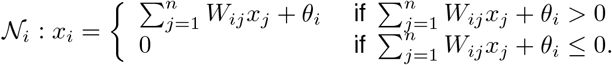

Figure 2C illustrates the hysteresis mechanism for the subnetwork in panel B. In the first subpanel (top left), the nullcline 𝒩_1_ is fully shown. The other nullclines 𝒩_*i*_ resemble 𝒩_1_, each consisting of a slanted part when *x*_*i*_ *>* 0 and a segment of a coordinate hyperplane with *x*_*i*_ = 0. The network has two stable fixed points, each lying at an intersection of nullclines. We restrict attention to the horizontal slices *x*_3_ = *x*_4_ = 0 and *x*_3_ = *x*_4_ = *a* where the fixed points lie. Initially, the network is settled in the (*a, a*, 0, 0) fixed point, where 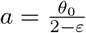 (top right subpanel). When the pulse comes, the value of *θ*(*t*) changes, causing the nullclines to slide upward (bottom left subpanel). As a result, (*a, a*, 0, 0) is no longer a fixed point, and the trajectory moves upward (*dx*_*i*_*/dt >* 0 for all *i*). When the pulse ends, the nullclines return to their initial positions. However, the trajectory does not return to the original (*a, a*, 0, 0) fixed point. Instead, it settles into the second fixed point, (0, 0, *a, a*), supported on the 3 ↔ 4 clique.

In the full counter (panel A), there are six stable fixed points, and the selected fixed point reflects the history of the number of pulses that have transpired. A natural question is: can we adapt this construction to a signed counter, where “−” and “+” pulses shift the counter left and right?

#### Signed TLN counter

To create a counter that can track both “− “and “+” pulse inputs, we begin from the TLN counter architecture but reverse the direction of the bottom cycle through the even numbered nodes. For a + input, a pulse is sent to the nodes in the top cycle only (odd numbered nodes); these are connected in a way that will advance the counter activity to the right. For a − input, a pulse is sent to nodes in the bottom cycle only (even nodes), which causes the counter activity to shift left. This allows the network to track relative number of “− “v.s. “+” inputs. Interestingly, the signed counter is more sensitive to pulses than the original TLN counter and is more likely to advance through multiple cliques at once in an uncontrolled “roulette” style cycling (see Supplementary Materials section A.2). However, when we introduce a brief refractory period after each pulse, the signed counter performs as desired, only advancing by a single clique at a time.

The core motif analysis in this case is the same, as the cliques remain target-free and the two cycles again survive by Theorem 1. Computationally, we also found | FP(*G*)| = 369 and

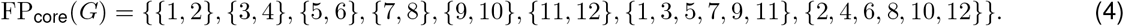

Again the predicted behavior of the network is confirmed in simulations, as seen in the rate curves of Figure 2D. The dynamics of this new counter are analogous to those of the previous counter, except now the count can be decreased. When the network receives an input pulse to the plus neurons (top cycle, black pulses in Figure 2D), the activity slides to the right. If it receives an input pulse to the minus neurons (bottom cycle, red pulses in Figure 2D), the activity slides to the left. That is, the activity travels in the direction of the cycle that received the pulse. This network can not only keep a count, but also store net displacement, position on a line, or relative number of left and right cues.

The sequences of fixed point attractors in both the signed and unsigned TLN counters were obtained by chaining together core motifs (2-cliques) that were guaranteed to yield stable fixed points. Similar ideas can be used to construct a more complex counter network where the individual states representing the count are dynamic attractors [60]. In addition, even though all the networks presented in this section keep a count modulo 6 (= number of chained motifs)—which could be useful when estimating time intervals or angles—this can be generalized to an arbitrary (but finite) number of motifs, allowing the network to keep a memory of an arbitrarily large count. Thus, the number of possible states approaches infinity as the number of links in the chain approaches infinity as well. Importantly, the networks scale linearly with the number of possible states, requiring two neurons per extra state. In contrast to traditional integrators, which are highly sensitive to parameter tuning, the models we present here are robust to both parameter variations and noise. Simulations show a wide range of parameters (*ε, δ, θ*) where the counters effectively track the number of input pulses received (see Supplementary Figure S2). However, performance can degrade in certain parameter regimes (e.g., rouletting behavior, stalls in a clique, or counts in multiples of two). These behaviors were studied for varying values of *ε, δ*, pulse intensity, and duration in Supplementary Figure S2.

The TLN counters are also robust to noise in the connectivity matrices *W* and in the inputs *θ*(*t*) (Figure 2E). In contrast, traditional neural integrators often require fine-tuning and can be highly sensitive to parameter variations. Thus, these models offer a simple yet robust alternative, responding well to a wide range of input strengths. A similar simple structure has been observed in flies, which can perform neural integration with remarkably small circuits [14]. The limits of this robustness is further explored in Supplementary Figure S3, where we show failure cases such as excessive sliding, getting stuck, or both.

### 2.3 Coexistence of dynamic attractors for five distinct quadruped gaits

A classical example in neuroscience that can be modeled with coexistence of dynamic attractors is animal locomotion [30, 62, 63]. Modeling the different modes of locomotion has been challenging because of the heavy overlap in units controlling limbs between different gaits. In addition, engineering the coexistence of several dynamic attractors is difficult because basins of attraction of nonlinear systems are often too intricate to reliably access each attractor [54, 55]. Most models overcome this difficulty by requiring changes in network parameters, such as synaptic weights, in order to transition between different gaits [30, 64, 63]. Here, we present a CTLN capable of reproducing five coexistent quadruped gaits: bound, pace, trot, walk and pronk; without any change of parameters needed to access different gaits. These gaits truly coexist as distinct limit cycle attractors in the network where different gaits are accessed via different initial conditions, or by stimulation of a specific neuron involved in the gait, as shown in Figure 1D3.

#### Math interlude: cyclic unions as pattern generators

One important architectural family of CTLNs that gives rise to patterned activations is the *cyclic union*, defined in [45] as follows: given a set of component subgraphs 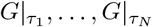, on subsets of nodes *τ*_1_, …, *τ*_*N*_, the *cyclic union* of 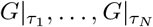 is constructed by connecting these subgraphs in a cyclic fashion so that there are edges forward from every node in *τ*_*i*_ to every node in *τ*_*i*+1_ (cyclically identifying *τ*_*N*_ with *τ*_0_), and there are no other edges between components (Fig. 3A). Remarkably, for cyclic unions it is possible to completely determine the fixed point supports of the full network in terms of the component fixed point supports:

**Figure 3:**
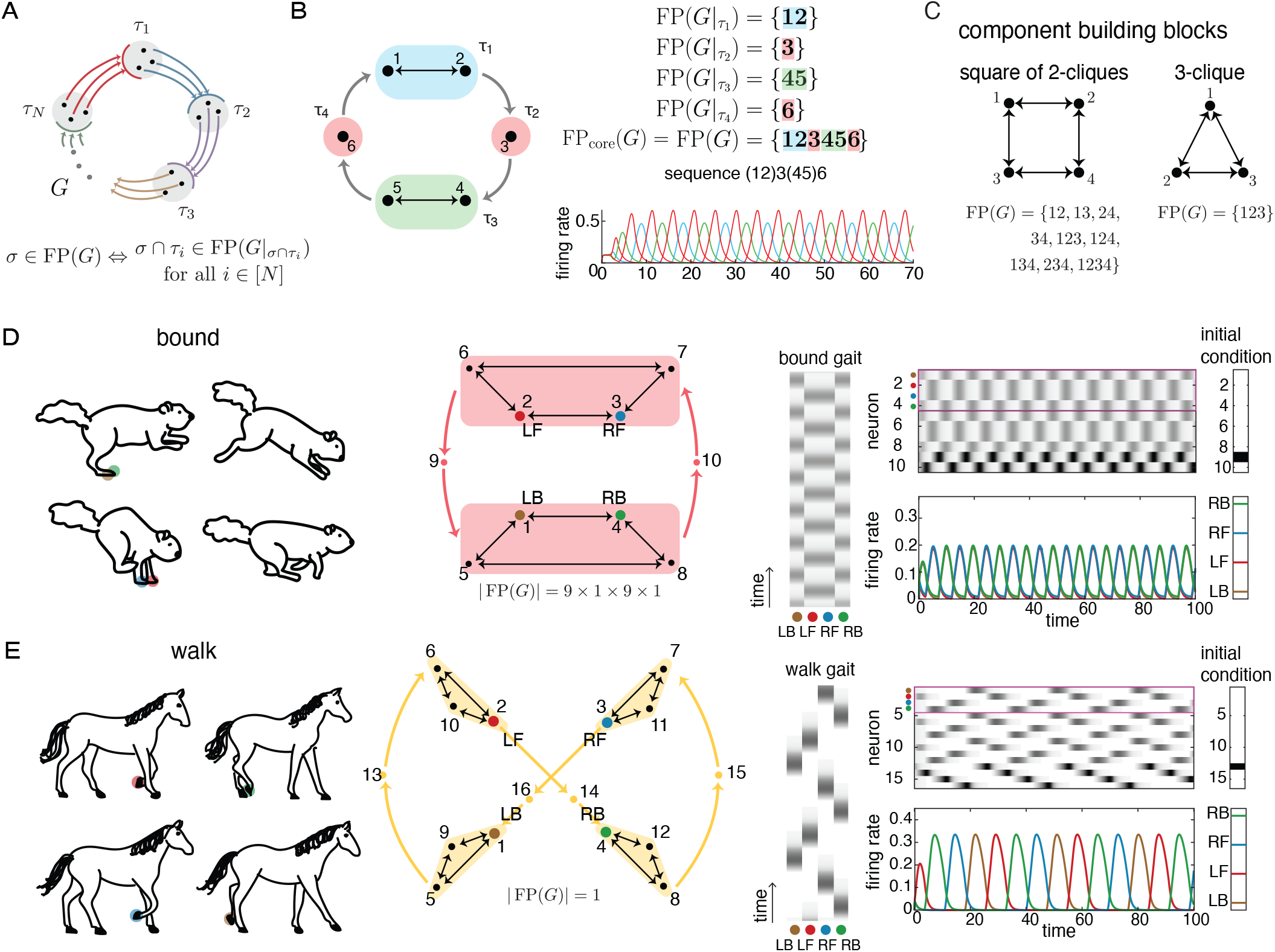
Design of quadruped gaits. (A) Cyclic union theorem. (B) Example of a cyclic union with two 2-cliques and two single nodes. FP(*G*) of the whole graph is made up of pieces taken from each 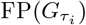, colored-coded by component. Notation: {*i*_1_, …, *i*_*k*_} is simplified as *i*_1_ …*i*_*k*_ (e.g., 12 = {1, 2}). Neurons 1,2,4, and 5 represent limbs, producing a bound gait. The firing rates were obtained using parameter values *ε* = 0.51, *δ* = 1.76, *θ* = 1. (C) The basic building blocks for each isolated gait, and their FP(*G*) sets. (D) Souslik’s bound, adapted from [61]. Colored circles represent node-limb assignment. Red shaded blocks group nodes in the same component. Red arrows represent edges from/to all nodes in a single component. Nodes 1 through 4 control the limbs (LB, LF, RF, RB). Nodes 5 to 10 are auxiliary nodes. The first four rows of the grayscale, outlined in pink and reproduced vertically, show the activity of the limbs. The network was initialized at *x*_9_(0) = 0.1 and all other neurons at 0. (E) Horse’s walk, adapted from [61]. Colored circles represent node-limb assignment. Yellow shaded triangles group nodes in the same component. Yellow arrows represent edges from/to all nodes in a single component. Nodes 1 through 4 control the limbs (LB, LF, RF, RB). Nodes 5 to 16 are auxiliary nodes. The first four rows of the grayscale, outlined in pink and reproduced vertically, show the activity of the limbs. The network was initialized at *x*_14_(0) = 0.1 and all other neurons at 0.

**Theorem 2** (cyclic unions, Theorem 13 in [45]). *Let G be a cyclic union of component subgraphs* 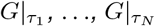. *For any σ ⊆* [*n*], *let* 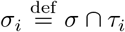. *Then*

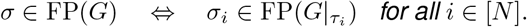

*Moreover*,

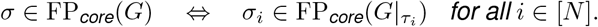

Note that, in particular, *σ*_*i*_ ≠ *∅* for all *I* ∈ [*N*] and so Theorem 2 guarantees that every fixed point of *G* hits every component. Since the core motifs of a cyclic union are found by taking a core motif from each component, we can patch together core motifs to follow cyclic activations using Theorem 2. For example, the quadruped gait bound is characterized by having both front limbs synchronized, both back limbs synchronized, and then successively cycling through these pairs, half a period off. This cycling makes it natural to group front limbs (nodes 1 and 2 in Figure 3B) in a single component, *τ*_1_, back limbs (nodes 4 and 5) in a single component, *τ*_3_, and to connect them through two auxiliary 1-node components, *τ*_2_, *τ*_4_, to ensure that the phase between groups of limbs is half a period off. Theorem 2 then says that FP(*G*) is made up of 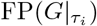 pieces, *i* ∈ [4]. Since each *τ*_*i*_ is a core motif (single nodes and 2-cliques), we obtain FP(*G*) = FP_core_(*G*), as detailed in Figure 3B. Notice that in the resulting attractor every component is active, and the neural activity flows through the components in cyclic order. If we take nodes 1, 2, 4, and 5 to represent limb nodes, then the network in Figure 3B successfully reproduces the activations seen in the bound gait.

#### Single gaits

In the previous example, bound was constructed using only six nodes. In what follows, we aim to construct a single network that encodes multiple gaits. However, having so few nodes per gait becomes impractical, as adding new gaits would significantly increase the number of edges between those nodes. This would cause the six-node network in Figure 3B to collapse into a clique, ultimately yielding a stable fixed point, as per Theorem 1. To address this, we propose a slightly different construction for bound, introducing a few extra auxiliary neurons per limb to mitigate this issue.

The resulting network is shown in Figure 3D, where we replace the previous 2-clique components with a square structure composed of four 2-cliques. The FP_core_(*G*) of this square consists of all the individual 2-cliques that form its sides, as illustrated in Figure 3C.

Theorem 2 now says that FP(*G*) of the bound gait network of Figure 3D is obtained by selecting a support 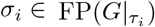 from each component *τ*_*i*_ and forming their union: *σ*_1_ *∪* {9} *∪ σ*_2_ *∪* {10}, where *σ*_1_ ∈ {{2, 3}, {2, 6}, {6, 7}, {3, 7}}and *σ*_2_ ∈ {{1, 4}, {1, 5}, {5, 8}, {4, 8}}. This includes, among others, the support {1, 2, 3, 4, 13, 14} corresponding to the bound attractor seen in the rate curves of Figure 3D. In an analogous way, it is possible to similarly construct pace (left limbs are synchronized, right limbs are synchronized, half a period off) and trot (diagonal limbs are synchronized, half a period off). The core motifs are similarly derived, using the “square” building block of Figure 3C, along with Theorem 2.

A slightly different construction is needed for the gait walk, where limbs move cyclically with a quarter period difference (Figure 3E). Again, we use Theorem 2 to group each limb in a component along with its auxiliary neurons and add four auxiliary neurons to ensure that the phase between limbs is a quarter period (Figure 3E). The building blocks are now copies of the 3-clique of Figure 3C. Since the building block is a core motif, the FP_core_(*G*) of the walk gait consists of a single element, formed by taking a core motif from each component in Figure 3E. Again, this design trick of adding extra auxiliary neurons is needed for the 5-gait network construction ahead. The resulting dynamics of this network are depicted in Figure 3E, and the network once again behaves as expected. The last gait our model includes is pronk, where all limbs are synchronized, requiring now a single component consisting of all four limbs together along with a pair of auxiliary nodes that control the frequency of the pronking movement.

All five networks described above are pictured in Figure 4A. We computationally derived the sizes of each of the FP(*G*) for each gait networks as designed in the previous paragraphs. These results, along with a summary of the previous discussion on their FP_core_(*G*) can be found in the Supplementary Table 2. Additionally, to verify that the core motifs accurately predict the attractors, we simulated each of the gait networks as previously constructed. The results are presented in Supplementary Figure S4D, which includes the grayscale and rate curves for all isolated gaits, along with the induced subgraphs for each gait.

**Figure 4:**
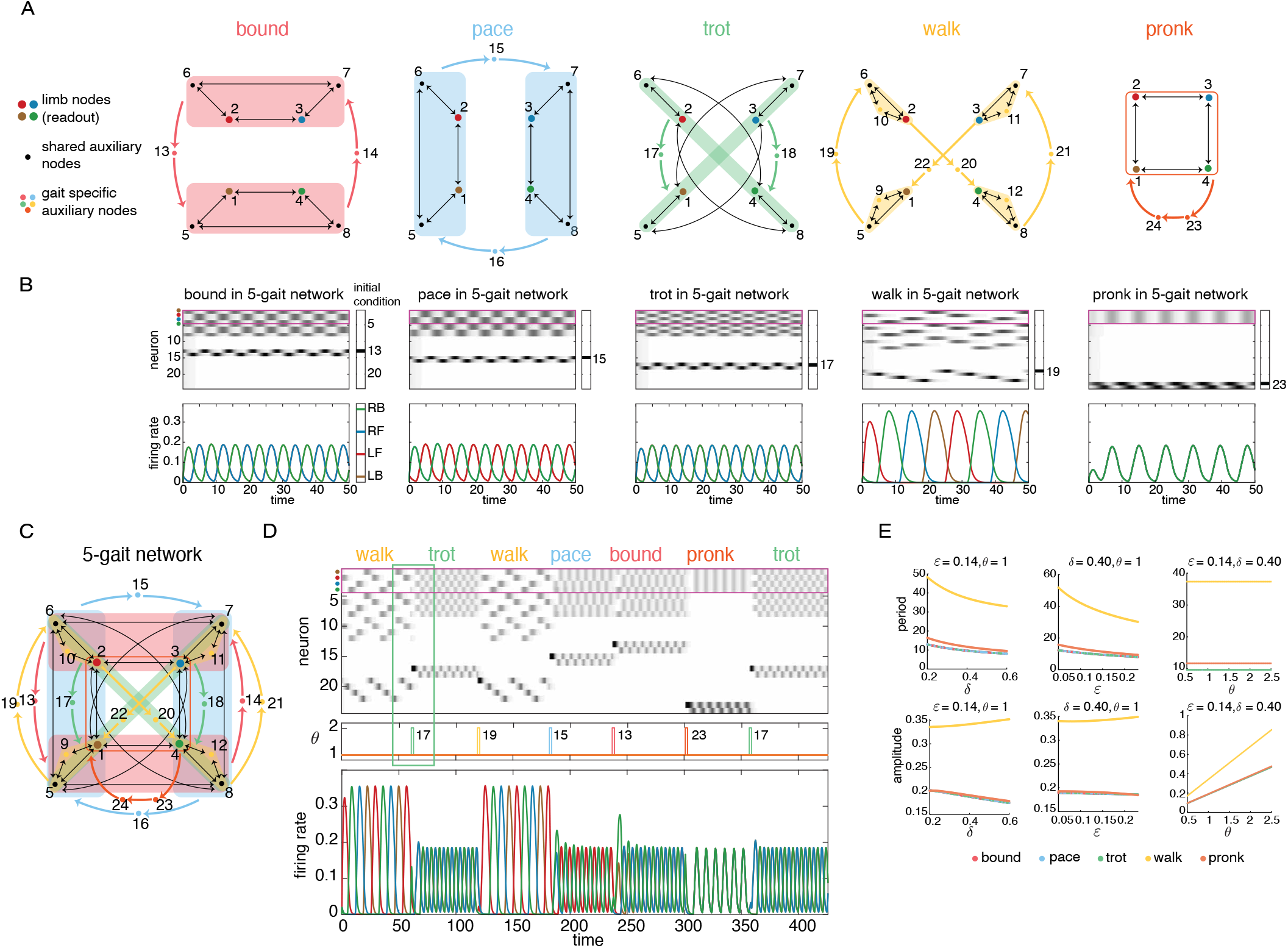
5-gait network and transitions. (A) Graphs for gaits constructed individually according to Theorem 2. Color-shaded areas group nodes in the same component. Colored arrows represent edges from/to all nodes in a single component. (B) Simulation of all 5 gaits within the network from panel C below. Initial conditions were set to zero for all neurons except for neuron *x*_*i*_ = 0.2, where *i* corresponds to the neuron specified in the panel next to each grayscale. The pink square highlights the activity of the readout neurons (limb nodes). (C) 5-gait network after gluing all individual networks from panel A (note that the networks in panel A are not the induced subgraphs of the 5-gait network). (D) Gait transitions via *θ* pulses. The neuron being stimulated with the pulse in the middle plot is written and colored according to which gait it corresponds to. The duration of the pulse is 2 time units. The activity quickly transitions to the attractor representing the corresponding gait. The order in which neurons are stimulated does not matter. The green box highlights the activity around the first pulse. (E) Amplitude and period of the gaits in the 5-gait network as controlled by *ε, δ, θ*. Gaits are color coded as in panel D. Values were found using the last half (once the attractor has settled) of a simulation with running time *T* = 500.

#### 5-gait network

To construct the 5-gait network, we glued together the networks from Figure 4A by identifying common neurons (1 through 8) along with all induced edges between them. The resulting glued network is shown in Figure 4C, this is the 5-gait network. More precisely, this gluing identifies all copies of nodes 1 to 8 from each isolated gait to a single node in the final network, along with their induced edges. This operation does not correspond exactly to a standard graph operation; rather, it is a topological gluing, where the identification rules are determined by the vertex numbers and the edges between them.

As mentioned before, the inclusion of several auxiliary nodes was necessary to merge the five networks of Figure 4A into a single one. Indeed, nodes 1 to 4 are shared by all gaits and represent the limbs: 1. LB - left back, 2. LF - left front, 3. RF - right front, 4. RB - right back. Nodes 5 to 8 are auxiliary (not representing limbs) and are shared by bound, pace and trot. The rest of the nodes are gait-specific, meaning that they are uniquely associated to a single gait and therefore only active when such gait is active, and are off otherwise. These nodes play the important roles of (1) keeping the phase between foot steps in each gait and (2) activating a specific gait by stimulating one of them. These auxiliary nodes are: nodes 9 to 12, specific to walk; and nodes 13 to 23 which are specific to a unique gait, as color coded in Figure 4A.

Although the complex structure of the glued network makes it hard to analytically derive the FP(*G*), we computationally found its core motifs (Supplementary Materials Table 2). Interestingly, none of these core motifs precisely correspond to the gait attractors, and yet all gait attractors are in fact preserved in the glued network, as seen in Figure 4B. The corresponding grayscale graphs and rate curves for each gait within the glued network are plotted below the respective graph in Figure 4A. Furthermore, the dynamics of each gait, when embedded in the glued network, are remarkably similar to those of the isolated networks. This is further illustrated in Supplementary Figure S4D, which shows the grayscale plots and rate curves for all isolated gaits, along with the induced subgraphs for each gait.

To see that each gait is accessible via different initial conditions, observe in Figure 4B that when all neurons, but neuron 13, are initialized to zero, the system evolves into the bound gait. This is because neuron 13 corresponds *uniquely* to bound. Similarly, when the network is initialized on neuron 15, the dynamics go into pace. When it is initialized on neuron 17, the dynamics go into trot. And so on. Recall that walk has two different sets of gait-specific neurons (9 - 12 and 19 - 22), and both kinds will send the network into the walk gait. Hence, our 5-gait network successfully encodes these coexistent dynamic attractors that have several overlapping active nodes (nodes 1 to 8), with no interference between attractors. Moreover, the attractors are not overly sensitive to the value of initial conditions, as long as the highest firing neuron in the initial condition is one that is uniquely associated with the desired gait.

Moreover, since all gaits coexist as distinct limit cycle attractors in the network, each accessible via different initial conditions, it is also possible to smoothly transition between gaits by means of a targeted external pulse *θ*. Specifically, it suffices to stimulate an appropriate gait-specific auxiliary neuron with a pulse. The network quickly settles into the dynamic attractor corresponding to the gait of the auxiliary neuron stimulated, as seen in Figure 4D. There, pulses are sequentially sent to neurons 17, 19, 15, 13, 23 and 17, producing the expected transitions between gaits. Any sequence of pulses sent to gait-specific auxiliary neurons will produce the respective transitions, regardless of the desired ordering of the gaits.

Finally, we evaluated the network’s performance under several parameter values, and show how the amplitude and period of the firing rate curve for each gait can be modulated by these parameters, as shown in Figure 4E. Increasing *ε* and *δ* resulted in a decrease in the period, while the period remained flat when varying *θ*, meaning the timing was unchanged but the intensity of firing increased. Notably, the walk gait showed an increase in amplitude, while the other gaits exhibited reduced amplitude as *δ* increased. For additional examples of how varying *ε, δ*, and *θ* affect the attractors (e.g., preserved symmetry, loss of symmetry, or switching to a different attractor), see Supplementary Figure S5. Details on how the period and amplitude of oscillations in firing rates change with these parameters, and how these values were obtained, are provided in Supplementary Figure S6.

Additionally, we evaluated the network’s performance under noise added to both the connectivity matrix *W* and the input signal *θ*. Bound, pace, and trot gaits remained stable with up to 2% noise but began to break down at 4%. In contrast, walk and pronk gaits were robust to noise levels as high as 10% and beyond. Supplementary Figure S7 further explores how gaits behave under varying levels of noise in both *W* and *θ*. While all gaits survived with up to 2% noise, others withstood higher levels but exhibited distorted behavior; the attractors remained, but their dynamics were less stable. This figure also examines transitions between gaits under noisy *W* and *θ*, showing that while transitions were disrupted at higher noise levels, they remained relatively stable at 1% noise.

### 2.4 Sequential control of quadruped gaits

In this section, we design a network that encodes a sequence of gaits *internally*, in contrast to the setup in Figure 4D, where each gait was selected by targeted input pulses, meaning any sequence was determined *externally*. To achieve internal encoding, we use fusion attractors, which combine the activity of the counter network from Figure 2A with the 5-gait network from Figure 4C. The objective is to create a network that, given identical inputs (with externally controlled input timing), outputs a predetermined sequence of gaits (or any set of dynamic attractors), as illustrated in Figure 1D4.

Fusion attractors are formed by combining stable fixed points from the counter network with dynamic attractors representing gaits from the 5-gait network, as illustrated in Figure 5A. This combination of stable fixed point with gait attractors results in a total of 5 · *m* fusion attractors, coming from 5 possible gaits and *m* possible fixed points from the counter.

**Figure 5:**
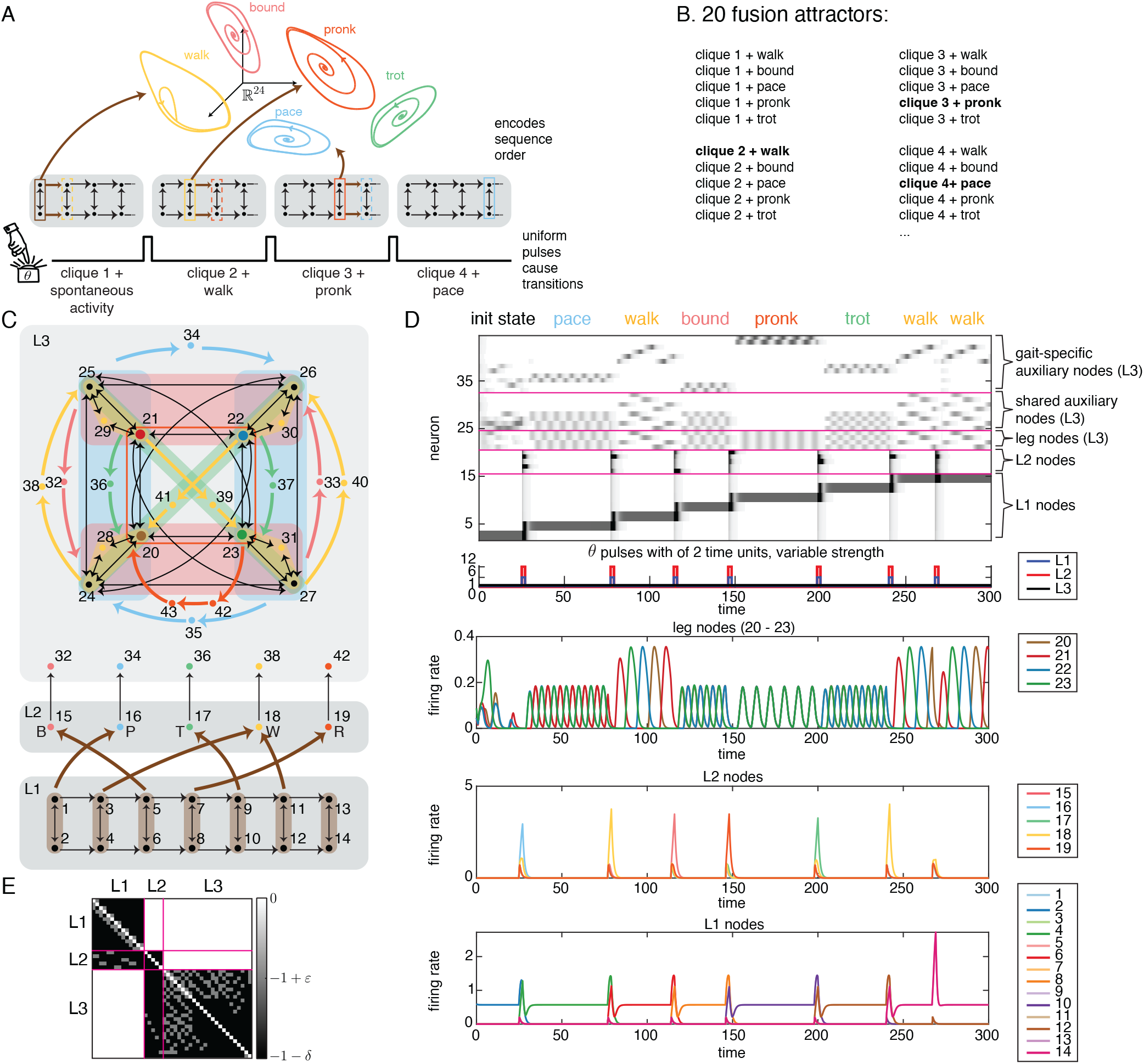
Sequential control of quadruped gaits. (A) Network design. Multiple attractors coexist in ℝ^24^ (state space of the quadruped gaits network), accessed via uniform global pulses (*θ*-pulses) sent to the TLN counter network (gray). Each pulse triggers the activity to advance following the brown arrows. The sequence order is encoded by the brown connections, separately from the attractors. The active clique is highlighted with a solid box, and the clique to be activated is highlighted with a dashed box, color coded by the gait it will be coactivated with. (B) List of fusion attractors that are possible from the gait attractors and counter network pictured in A. Out of the 5 · 4 possible attractors, 3 of them are selected by the prescription of the brown arrows of A. Selected attractors in bold. (C) Layered network for sequential control of quadruped gaits. L1 (unsigned counter) encodes the sequence steps. L2 (independent set) acts as relay layer, each node connects to an attractor-specific node in L3 (5-gait network). Attractor-specific nodes corresponding to different gaits are redrawn at the bottom for clarity. Every node within the shaded block sends edges to where the thick arrow points. (D) Grayscale, pulses and rate curves. On top of the grayscale is the order in which gaits are accessed, matching those specified by the brown arrows in C. Only neurons in L1 and L2 receive pulses. L2 does not receive any external input outside of the pulse times. There are four different *θ* values pictured. Neurons in L1 have a baseline *θ* = 1 and pulse *θ* = 6. Neurons in L2 have a baseline *θ* = 0 and pulse *θ* = 12. Neurons in L3 have a constant *θ* = 1. Note that transitions between cliques in L1 are communicated effectively as pulses to L3 via L2. The pairs of attractors that are fused are the ones that get co-activated by the prior clique after a pulse. For example, starting in clique {1, 2}, when the pulse comes, both clique {3,}4 and the pace gait get stimulated, producing {3, 4} -P as the first fusion attractor. (E) *W* matrix for the network in C.

Figure 5A shows an example where the counter consists of four 2-cliques (stable fixed points). Out of the possible 5 · 4 = 20 fusion attractors listed in Figure 5B, only four are activated (bold). These are determined by the thick brown arrows—every gait-clique pairing that receives brown arrows from the same clique will be activated as a fusion attractor. For instance, the first fusion attractor triggered in Figure 5A is clique 2 + walk, since both receive edges from the same (previous) clique. Note that the cliques are always stepped through in order, but their pairing can be with any gait, dictated by the connectivity between the previous clique and the gait network. As a result, gaits can be executed in any pre-specified order. Also note that the external pulse is what drives the sequence forward.

In practice, directly connecting the counter network to the 5-gait network caused interference, where the fixed-point attractors of the counter disrupted the dynamic attractors of the quadruped gaits network. To resolve this, we introduced an intermediate relay layer (see Figure 5C) that is inactive except during the pulse transitions. The attractors of this 3-layer network include 7 · 5 = 35 fusion attractors, each consisting of one of the 7 stable fixed points from the cliques in layer L1 together with one of the 5 gaits from layer L3. While all 35 fusions are technically attractors of the network, in practice, only 6 of them are typically accessed; specifically, the sequence of 6 that are encoded in the network via the connections between L1, L2, and L3 (see thick brown arrows in Figure 5C).

From the prescription of brown arrows in Figure 5C, the network encodes the sequence: clique 2 + pace, clique 3 + walk, clique 4 + bound, clique 5 + pronk, clique 6 + trot, clique 7 + walk, accessed in that order by identical pulses. While gaits can be repeated in the sequence, each fusion attractor is different from previous repetitions of that gait. For instance, clique 3 + walk and clique 7 + walk both output the same gait (walk), but they correspond to different fusion attractors. This difference in attractors is key to remembering location in the sequence, as L1 keeps track of which step in the sequence is active.

Fusion attractors that are active are easily identifiable in the grayscale of Figure 5D. Recall the fusion attractor pairings are determined by the thick brown arrows—everything receiving an brown arrow from the same clique will be fused together. For example, pronk fuses with clique 5 as both pronk (via node 19) and clique 5 receive inputs from clique 4.

Also, note that the input pulses in Figure 5D, which target layers L1 and L2, are identical and uniform within each layer. Importantly, layer 2 receives no external *θ* input outside of pulse times; specifically *θ*_*L*1_ = *θ*_*L*2_ = 1 outside of pulse times, while *θ*_*L*3_ = 0. The network’s output corresponds to the activity of the readout neurons, or limb neurons, which directly encode the gaits.

Finally, as seen in Figure 5E, this network is not technically a CTLN, as the off-diagonal values are not exclusively −1 + *ε* or −1 − *δ*. Non-edges within a layer and non-edges from one layer to the next are valued at −1− *δ*, while all other non-edges are set to 0. Consequently, the matrix is sparse, unlike CTLNs, with many zeros in addition to strong and weak inhibition.

### 2.5 Why does this work? The theory behind the intuition

The mechanism behind the network in the previous section is actually provably simple. It involves two distinct networks that are transiently coupled through a relay layer, as seen in Figure 6. To formalize this, we need to establish the existence of fusion attractors in networks with this structure. Specifically, these are networks with one layer encoding only stable fixed points, an intermediate relay layer, and another layer encoding dynamic attractors. In fact, the structure of the network in the previous section can be further relaxed.

**Figure 6:**
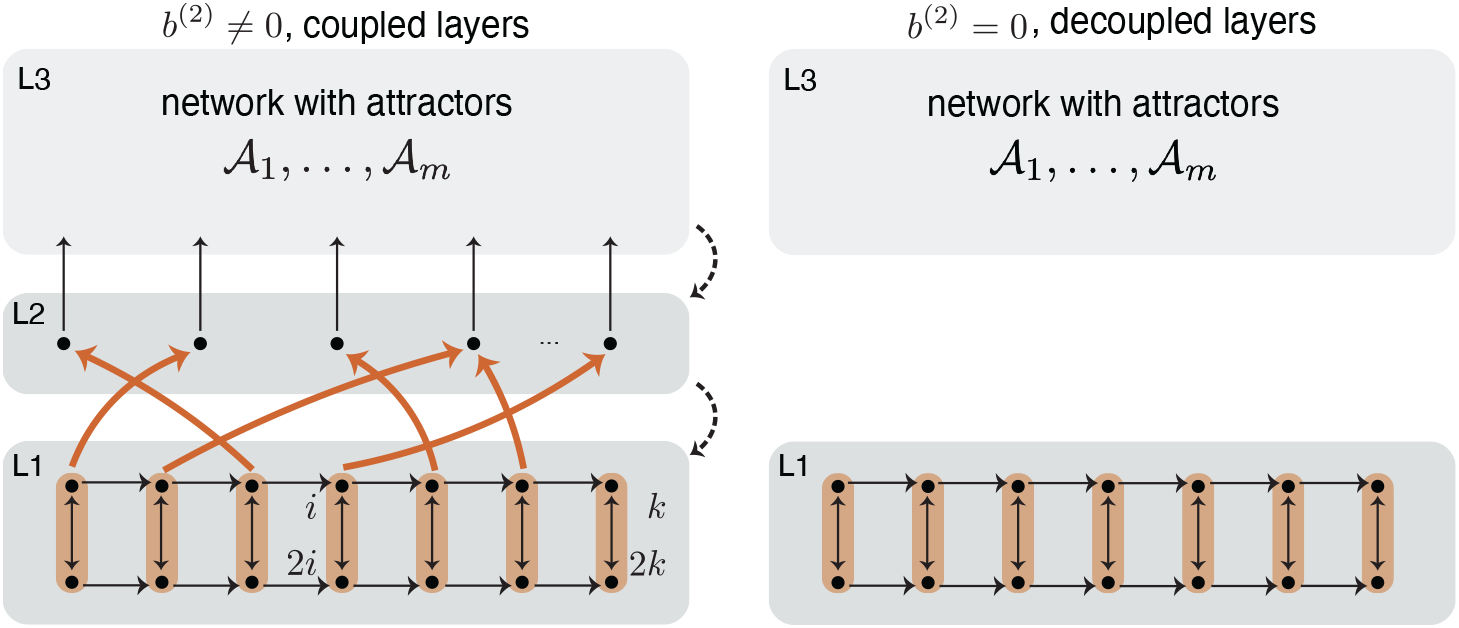
Fusion attractors obtained via decoupled layers. (A) General structure of network for Theorem 3. L1 is a network with *k* stable fixed point attractors coming from the cliques. L2 is an independent set of *m* nodes. L3 is an arbitrary network with *m* arbitrary attractors. The only connections allowed by the theorem are between consecutive layers (black, orange and dashed edges). (B) The resulting network after the external input is turned off for layer 2 (*b*^(2)^ = 0).

**Theorem 3**. *Consider a TLN* (*W, b*) *where*

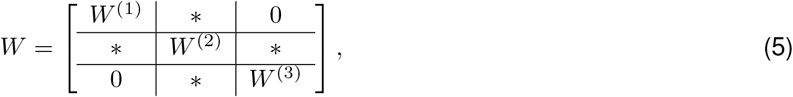

*and*

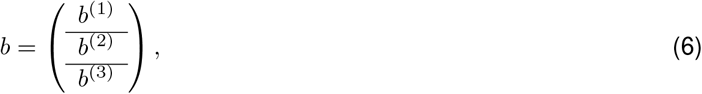

*and suppose that* (*W* ^(1)^, *b*^(1)^) *is a network with k stable fixed points and no other attractors;* (*W* ^(2)^, *b*^(2)^) *is an independent set on* ℓ *nodes; and* (*W* ^(3)^, *b*^(3)^) *is an arbitrary recurrent network with attractors* 𝒜_1_, …, 𝒜_*m*_.

*Then, if b*^(2)^ *≤* 0, *the attractors of the network* (*W, b*) *include the m* · *k fusion attractors consisting of one stable fixed point attractor coming from* (*W* ^(1)^, *b*^(1)^) *fused with one A*_*j*_ *attractor coming from* (*W* ^(3)^, *b*^(3)^).

The proof of this theorem is provided in Supplementary Materials Section A.4. This theorem specifically shows that the role of L2 is to transiently couple layers L1 and L3 during the pulse. Importantly, the theorem allows us to select among the attractors by initializing the system in the desired attractor in L1 and L3. Since the attractors are robust, the influence of L2 remains transient and minimal. Similar results have been established for other architectures involving CTLN layers, which can be found in [60].

Note that for the specific case of L1 being the counter network in Figure 6 we have a stable fixed point for each clique, by Theorem 1. Because the cycles are absent in this modified counter network, there are no additional core fixed points. This leads us to conjecture that the only attractors in the L1 network of Figure 6 are the stable fixed points associated with the cliques. Combining this with Theorem 3, we have the following conjecture:

**Conjecture 4**. *The network in Figure 6 has exactly m* · *k fusion attractors given by clique i + A*_*j*_ *for i* ∈ [*k*], *j* ∈ [*m*]. This suggests that designing networks that step through attractors using this method should be relatively straightforward. Another example using a different set of dynamic attractors corresponding to the swimming directions of a marine mollusk can be found in [60].

## 3 Discussion

Our aim has been to unify models of rhythmic activity, traditionally modeled using coupled oscillators, with models of associative memory networks, traditionally modeled using attractor neural networks. This led us to choose the TLN framework, as it can produce a wide variety of dynamic behaviors, including coexistence of multiple static and dynamic patterns. Indeed, as we demonstrated in this work, the dynamic variety of TLNs makes them well-suited for modeling a range of neural functions. While these models are not intended as a direct biological analogs, they suggest mechanisms that could be tested in both neural and robotic systems. Indeed, locomotive patterns can be used as priors for learning, significantly accelerating adaptation in artificial systems [65].

First, we designed a network that encodes a count of external input pulses using multiple stable fixed points. Although this is similar in spirit to Hopfield networks, our network is not symmetric and no energy function exists. As a result, it can exhibit hysteresis, a property that enables the network dynamics to “remember” prior states. This hysteresis mechanism is precisely what allows the network to increase or decrease the count by moving through a series of attractor states in response to external inputs. This model provides a robust and simple alternative to traditional neural integrators and evidence accumulators [56].

The same TLN framework also allowed us to encode multiple dynamic attractors, with prescribed patterns of activation, in a controlled and deliberate way. We successfully encoded five distinct quadruped gaits—bound, pace, trot, walk, and pronk—within the same network. Remarkably, the four nodes controlling the limbs, which are common to all gaits, fire in distinct patterns depending on the active gait attractor. Additionally, rapid transitions between gaits are easily achieved by applying external targeted pulses.

Finally, by combining these two approaches, we applied the concept of fusion attractors (Fig. 1C) to create a network that can step through a sequence of quadruped gaits. In this case, the fixed point counter steps through a sequence of static attractors using external pulses, which in turn activates one of the dynamic patterns of the quadruped gait network. By integrating these two mechanisms, we translated a sequence of pulses into a sequence of gait activations.

This final construction preserves the key properties of the individual components. From the counter, we have an external controller that induces transitions between stored attractors. From the quadruped gaits, we have several patterns available to select from. From the conjunction, we get the freedom to choose the sequence order. All of this is integrated by the fusion attractors, where the fixed-point component remains unchanged, but the gait paired with each fixed point depends on the network’s connectivity. In other words, while the counter determines which fixed point is active, the network structure dictates which gait is associated with it. In this way, we encode the sequence of gaits within the network while maintaining external control over the timing of transitions, much like the choreography of dance moves. In summary, three benefits arise from the design of this model:

1. **Sequential information is internally encoded**. Since external pulses are sent to all neurons in layers L1 and L2, rather than to specific neurons controlling each gait (as in Section 2.3), the sequence is fully encoded within the network, and the pulses themselves carry no information about what the next step is. This contrasts with similar models for sequential control, which also rely on external inputs but typically require motif-specific inputs to activate each element in the sequence [66, 67].
2. **Sequential information is independently encoded from motor commands**. The order of the sequence is encoded in the connections from layer L1 to L2, while the dynamic attractors are encoded in L3. This disassociates the sequence order from the elements themselves, allowing the network to reuse elements by adding just a few extra neurons and connections. This efficient reuse of attractors is a remarkable feature, as dynamic attractors are usually very sensitive to interference [54, 55]. A similar reuse of dynamical motifs has been observed in recurrent networks performing multiple tasks, where motifs are shared across different computations [68].
3. **Timing of transitions is externally encoded**. Timing of the sequence is controlled by external pulses, and therefore a sequence’s execution can be sped up or slowed down, or the rhythm altered altogether, without interfering with attractors or sequence order. Moreover, there is no need to change synaptic time constants or physiological variables to alter the duration of each sequence step.

In summary, our unified attractor network framework not only allows rhythms and oscillations to be encoded alongside static memory patterns, but also provides simple constructions for modular architectures that enable the flexible binding and reuse of attractors in prescribed sequences.

## 4 Methods

### Network simulations (deterministic)

Each network was simulated using the following equations:

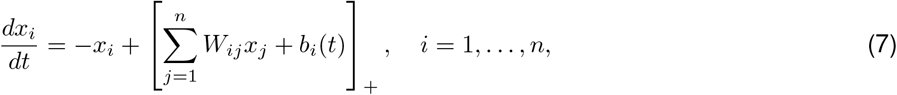

Here, *W* is obtained from the graph displayed in each figure, according to the prescription given by equation (2). The particular values of *ε* and *δ* vary per network, and can be found in Table 1.

**Table 1:** Parameters used in each network simulation. The initial condition *x*(0) is set with all neurons off, except those listed as ON, which are initialized to the value indicated under the “initial value” column.

| Network $G$ | $\varepsilon$ | $\delta$ | $\theta$ | ON neurons at $x(0)$ | initial value |
| --- | --- | --- | --- | --- | --- |
| fixed point attractor (Fig 1A) | 0.25 | 0.5 | 1 | 1, 2 | random |
| sequential attractor (Fig 1B) | 0.25 | 0.5 | 1 | 1, 2 | random |
| fusion attractor (Fig 1C) | 0.25 | 0.5 | 1 | 1, 2 | random |
| TLN counter (Fig 2A) | 0.25 | 0.5 | 1, 5 | 1, 2 | 0.3 |
| signed TLN counter (Fig 2D) | 0.25 | 0.5 | 0.95, 1, 2.5 | 1, 2 | 0.3 |
| bound (Fig 3D) | 0.25 | 0.5 | 1 | 9 | 0.1 |
| walk (Fig 3E) | 0.25 | 0.5 | 1 | 13 | 0.1 |
| all gaits in quadruped gaits network (Fig 4B) | 0.25 | 0.5 | 1 | 13, 15, 17, 19, 23 | 0.2 |
| gait transitions (Fig 4D) | 0.25 | 0.5 | 1, 2 | 19 | 0.3 |
| sequential control of gaits (Fig 5D) | 0.25 | 0.5 | 0, 1, 6, 12 | 1, 2 & 20 – 43 | 0.6, 0.6 & random |

**Table 2:**
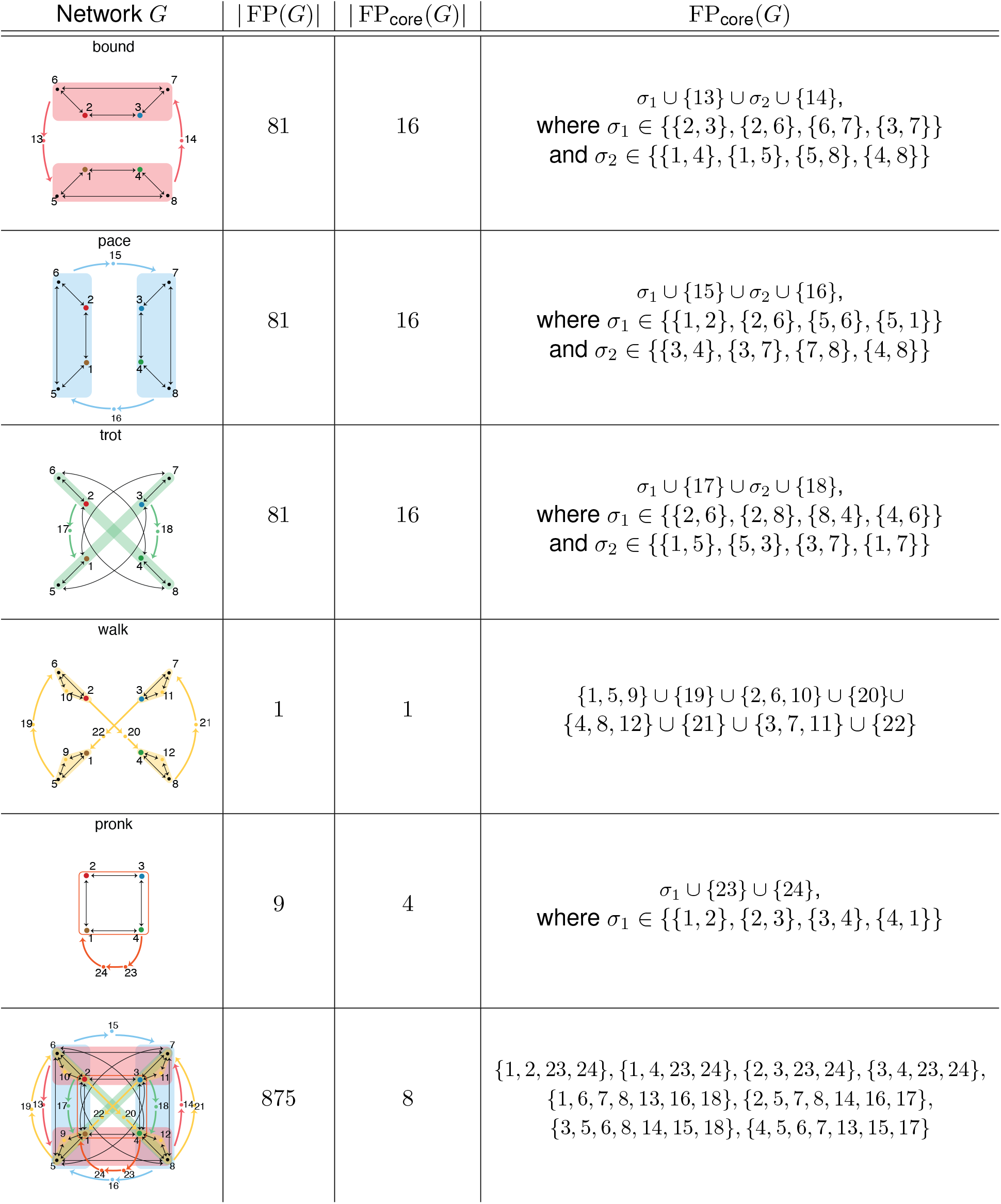
Gait networks’ FP_core_(*G*), for individual gaits and for entire network containing all gaits from Section 2.3.

| Network $G$ | $\vert FP(G)\vert $ | $\vert FP_{core}(G)\vert $ | $FP_{core}(G)$ |
| --- | --- | --- | --- |
| bound | 81 | 16 | $\sigma_1 \cup \{13\} \cup \sigma_2 \cup \{14\},$<br>where $\sigma_1 \in \{\{2, 3\}, \{2, 6\}, \{6, 7\}, \{3, 7\}\}$<br>and $\sigma_2 \in \{\{1, 4\}, \{1, 5\}, \{5, 8\}, \{4, 8\}\}$ |
| pace | 81 | 16 | $\sigma_1 \cup \{15\} \cup \sigma_2 \cup \{16\},$<br>where $\sigma_1 \in \{\{1, 2\}, \{2, 6\}, \{5, 6\}, \{5, 1\}\}$<br>and $\sigma_2 \in \{\{3, 4\}, \{3, 7\}, \{7, 8\}, \{4, 8\}\}$ |
| trot | 81 | 16 | $\sigma_1 \cup \{17\} \cup \sigma_2 \cup \{18\},$<br>where $\sigma_1 \in \{\{2, 6\}, \{2, 8\}, \{8, 4\}, \{4, 6\}\}$<br>and $\sigma_2 \in \{\{1, 5\}, \{5, 3\}, \{3, 7\}, \{1, 7\}\}$ |
| walk | 1 | 1 | $\{1, 5, 9\} \cup \{19\} \cup \{2, 6, 10\} \cup \{20\} \cup$<br>$\{4, 8, 12\} \cup \{21\} \cup \{3, 7, 11\} \cup \{22\}$ |
| pronk | 9 | 4 | $\sigma_1 \cup \{23\} \cup \{24\},$<br>where $\sigma_1 \in \{\{1, 2\}, \{2, 3\}, \{3, 4\}, \{4, 1\}\}$ |
| | 875 | 8 | $\{1, 2, 23, 24\}, \{1, 4, 23, 24\}, \{2, 3, 23, 24\}, \{3, 4, 23, 24\},$<br>$\{1, 6, 7, 8, 13, 16, 18\}, \{2, 5, 7, 8, 14, 16, 17\},$<br>$\{3, 5, 6, 8, 14, 15, 18\}, \{4, 5, 6, 7, 13, 15, 17\}$ |

The value of *b*_*i*_(*t*) also varies per network. The pulse values and the neurons pulsed were different for each network (as detailed below), but all of them had a baseline *b*_*i*_(*t*) = *b* = 1. Pulses could be either targeted to specific neurons (attractor-specific) or sent to all the neurons in the network (global). When pulses were targeted, the color of the pulse in the corresponding figure indicates which neurons received it. Values of the targeted pulses, along with its baseline, for each network are detailed below. The pulsed neurons and pulse times can be inferred from the pulse plot in each of the simulations, between the gray scales and the rate curves.

Each solution *x*_*i*_(*t*), for *i* = 1, …, *n*, was generated by solving the system of ODEs in equation (7), corresponding to each network as prescribed by the values in Table 1. The system was solved in MATLAB using a standard ODE solver (ode45). The initial conditions for all networks are: all neurons off (*x*_*i*_(0) = 0), except the ones specified in the “ON neurons” column of Table 1. Those in the ON neurons column were initialized to the value indicated in the “init” column, i.e. *x*_*j*_(0) = init for *j* an ON neuron.

The solution *x*_*i*_(*t*) is displayed in two different forms in each figure. On top there is a gray scale, where darker gray corresponds to higher firing rates. Below these are the pulses received by the network. And on the bottom are the rate curves of each *x*_*i*_(*t*).

## Acknowledgments

We would like to thank the members of our laboratory for their constructive feedback on the manuscript. Special thanks to Nicole Sanderson for her detailed comments. This work was supported by the grants NIH R01 EB022862, NSF DMS-1951165 and DMS-1951599. Part of this work was supported by the grant NSF DMS-1929284 while the authors were in residence at the Institute for Computational and Experimental Research in Mathematics (ICERM) in Providence, RI, during the Math + Neuroscience program in Fall 2023.

## A Supplementary materials

### A.1 E-I equivalence

In [53], it was shown that every CTLN is equivalent to an excitatory-inhibitory (E-I) network with the same connectivity graph and a single inhibitory node. When the timescale of inhibition is sufficiently fast, the E-I network matches the dynamics of the CTLN and the attractors appear to be identical as in Figure S1.

**Supplementary Figure S1:**
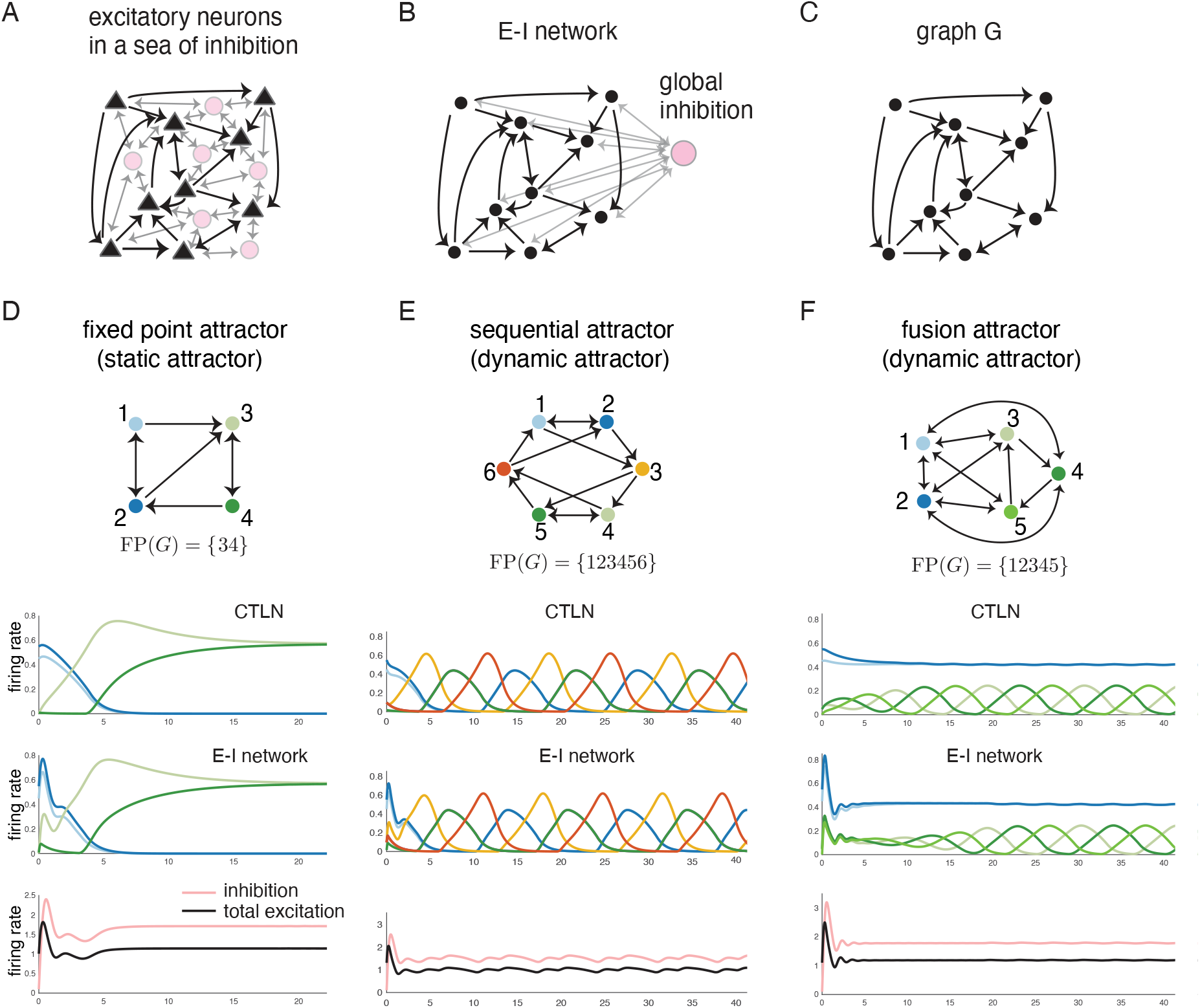
Examples of matching dynamics between CTLNs and equivalent excitatory-inhibitory (E-I) networks with the same connectivity graph between excitatory neurons, and a single inhibitory node [53]. (A) A network with excitatory neurons (black triangles) within a sea background of inhibition (pink circles). (B) E-I network with excitatory nodes (black) and a single inhibitory neuron (pink) accounting for all the background inhibition. (C) The connectivity graph for the CTLN and the equivalent E-I network consists of only the excitatory neurons and the connections between them. (D–F) Comparison of the dynamics between the CTLN model and the equivalent E-I model with the same connectivity graph. In all examples, the inhibition timescale is 5 times faster than that of excitation. Bottom panels show total inhibition (pink) and total excitation (black) for the E-I network

### A.2 TLN counters

#### Robustness of TLN counters

In general, CTLNs are expected to perform robustly, since much of their dynamic information is influenced by FP_core_(*G*) and we know this set is preserved across the legal range 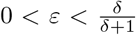 for model parameters *ε, δ* for certain networks.

To asses the performance of both counters across parameter space, we ran the simulations of Figure 2 using various pairs of (*ε, δ*) parameters, and various combinations of pulses strengths and durations. We found three different possible outcomes, exemplified in Figure S2A. We classified these according to whether the function is preserved (keeps the correct count), corrupted (counts in multiples of two) or lost (does not keep a consistent count) as *ε, δ, θ* vary.

**Supplementary Figure S2:**
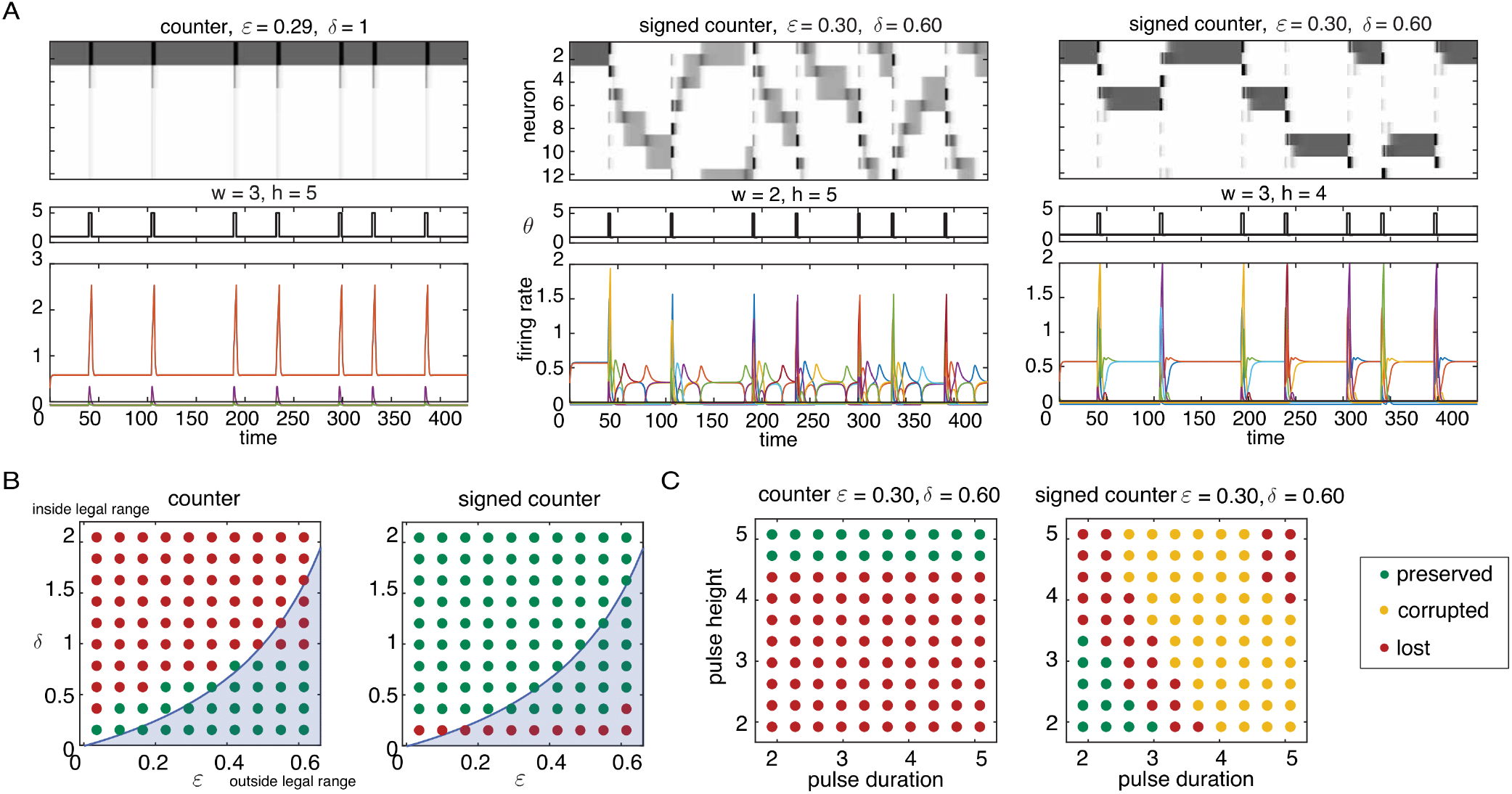
Good parameter grid for TLN counters. TLN counters’ behavior is classified according to whether the function is preserved (keeps the correct count, green dots), corrupted (counts in multiples of two, yellow dots) or lost (does not keep a consistent count, red dots) when *ε, δ, θ* vary. (A) Examples of what can go wrong. In the first plot the pulses do not move the counter from the current fixed point (function is lost), in the second plot the signed counters enter a “roulette” behavior (function is lost), and in the third plot the counter consistently slides two positions (function is corrupted). (B) Various counter behaviors for fixed values of baseline *θ* = 1, with pulses *θ* = 5 for counter, *θ* = 2.5 and *θ* = 0.95 for signed counter. *ε, δ* vary according to the axes. One dot correspond to one simulation of 7 pulses, with a running time of *T* = 428 time units for the counter and *T* = 435 time units for the signed counter. Shaded in blue are parameter pairs outside the legal range 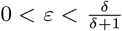. Green dot indicate no failures for the entire duration of the simulation. Any failure among the 7 pulses classifies as lost or corrupted function. (C) Various counter behaviors for a fixed values of *ε* = 0.30, *δ* = 0.60 and baseline *θ* = 1. Pulse height and duration vary according to the axes.

A brief refractory period after the pulse is needed for the signed counter to stop activity from sliding forward more than one clique, as otherwise the counter could enter the rouletting behavior of Figure S2, second panel.

The results for all pairs (*ε, δ*) of parameters and various combinations of pulses strengths and durations are recorded in the bottom row of Figure S2, where green indicates preserved, yellow indicates corrupted and red indicates lost performance. One dot corresponds to one simulation of 7 pulses, with a running time of *T* = 428 time units for the counter and *T* = 435 time units for the signed counter. We found that there is a good range of parameters where the counters behave as expected, successfully keeping the count of the number of input pulses received.

This robustness across CTLN parameter space is not surprising given that FP_core_(*G*) was preserved across the space. But will the performance of the counters survive added noise in the input and connections? To verify robustness to noise, we computed the proportion of failed transitions among *m* = 100 pulses in both fixed point counters. This was done by introducing varying percentages of noise into the *θ* input, varied every *dt* = 0.1 of a time unit, and into the connectivity matrix *W*, as specified in the axes of Figure S3. A failed transition is defined as any behavior that does not advance the counter a single clique forward/backwards, depending on the sign of the pulse. Figure S3 contains examples of all possible transition failures, as well as examples of successful transitions in noisy conditions, as detailed below. All simulations used to quantify the failures were done with *ε* = 0.25, *δ* = 0.5.

**Supplementary Figure S3:**
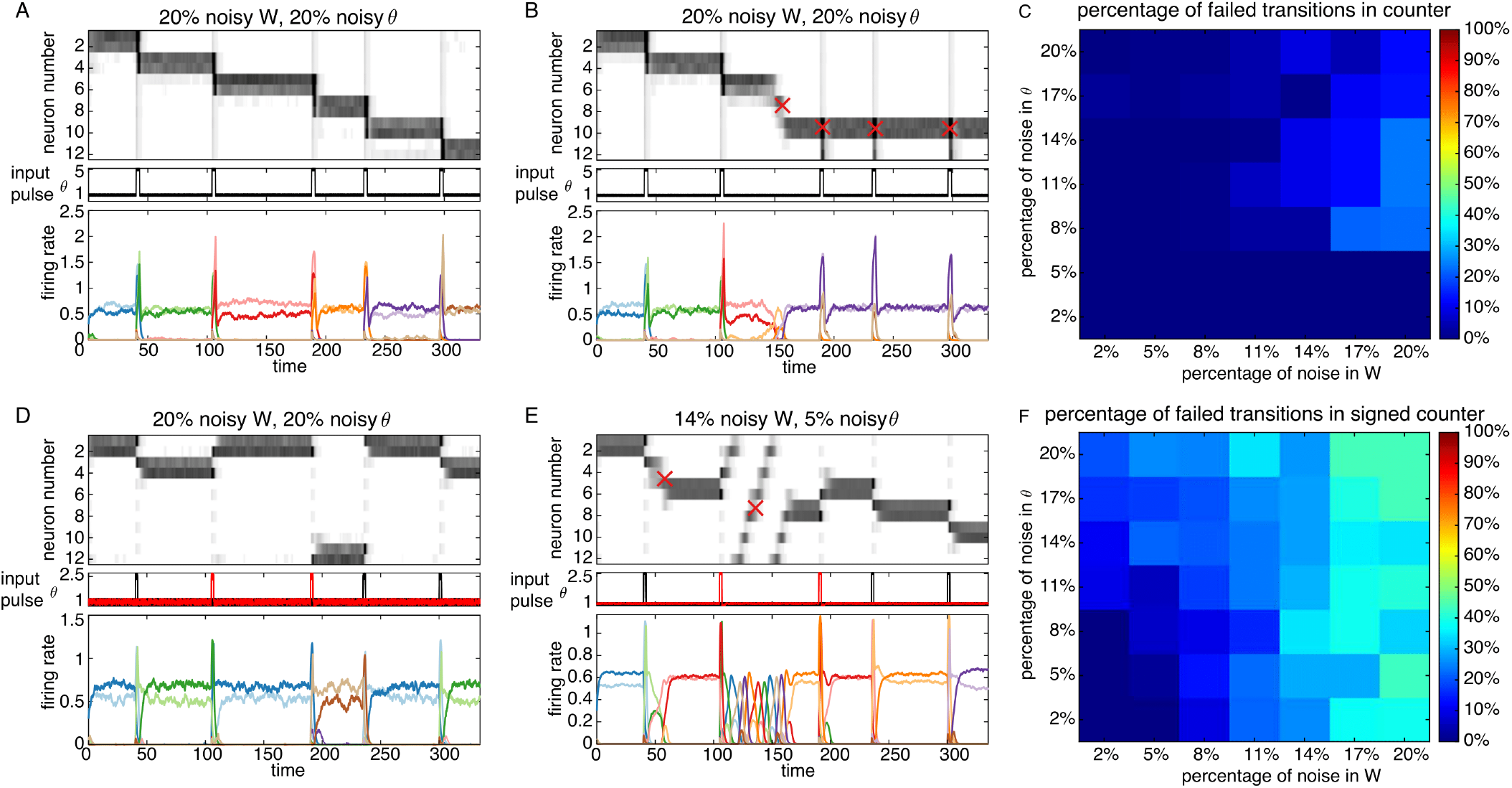
Noisy TLN counters. (A) Example of a 20% noisy counter that performs well. (B) Example of a noisy counter with four failures, marked with red crosses: first it slides too many cliques forward, and then it gets stuck in the same clique indefinitely. (C) Percentage of failed transitions for the counter in 100 trials, for several percentages of noise in *W* and *θ*. (D) Example of a 20% noisy signed counter that performs well. (E) Example of a noisy signed counter with two failures, marked with red crosses: first it slides too many cliques forward, and then it rolls around the counter until it settles in some arbitrary clique. (F) Percentage of failed transitions for the signed counter in 100 trials, for several percentages of noise in *W* and *θ*.

Panel A of Figure S3 is an example of a very noisy counter that still preforms well. Each pulse advances the counter a single clique forward, as expected. Panel B of Figure S3 shows a noisy counter with two types of failures: first the counter advances too many steps, and then it gets stuck in one of the cliques (altogether these would count as four failures in our analysis, even though there are probably just two defective cliques). Panel D of Figure S3 is an example of a very noisy signed counter that performs perfectly well. Panel E of Figure S3 is an example of things that can go wrong in the signed counter. The first failure advances the counter one more than expected, the second failure makes the counter go into a kind of roulette behavior until it stops at a random clique. We have also observed the signed counter getting stuck at some point, as in Figure S3B.

The percentages of noise in Figure S3 were calculated as follows: The noise in the external input *θ* is i.i.d random noise from the interval (−1, 1), and it was introduced to vary every 0.1 fraction of a unit time. The noise in the connectivity matrix *W* is obtained by perturbing the adjacency matrix *A* with i.i.d random noise from the interval (0, 1). More precisely, if denote by 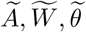 the noisy versions of *A, W, θ*; where *A* is the transpose of the standard graph-theoretic adjacency matrix of the graph *G* defining the network, then the noisy versions are:

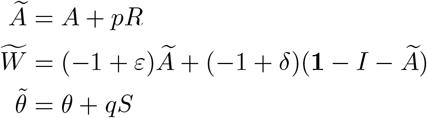

where *p, q* are the percentages of noise introduced, and *S, R* are random matrices the same size as *W, θ* with entries between (0, 1) and (−1, 1), respectively, and **1** is a matrix of 1’s the same size as *A*. This is equivalent to perturbing *W* by an amount proportional to length of the interval [−1 − *δ*, −1 + *ε*], that is:

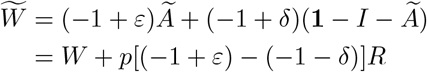

We choose to perturb the adjacency matrix instead for computational ease. For each percentage of noise pair (*p, q*), we ran 20 simulations, each one consisting of 5 (signed) pulses in the TLN (signed) counter, as exemplified in Figure S3. The results of counting over all these pulses are summarized in panels C and F of Figure S3. Each pixel represents the fraction of failed transitions for each pair (*p, q*). We found that discrete counters perform perfectly well up to 5% noise, establishing that perfect binary synapses are not necessary. When the noise limits are pushed beyond, counters lose stability and start to slide too many attractors, or to get stuck in one of the cliques. The (unsigned) counter, not surprisingly, proved to be a lot more robust, being barely affected by the amount of noise in *W*. Although not as robust, the signed counter accurately kept the count 50% of times under 20% noise.

### A.3 Quadruped gaits

**Supplementary Figure S4:**
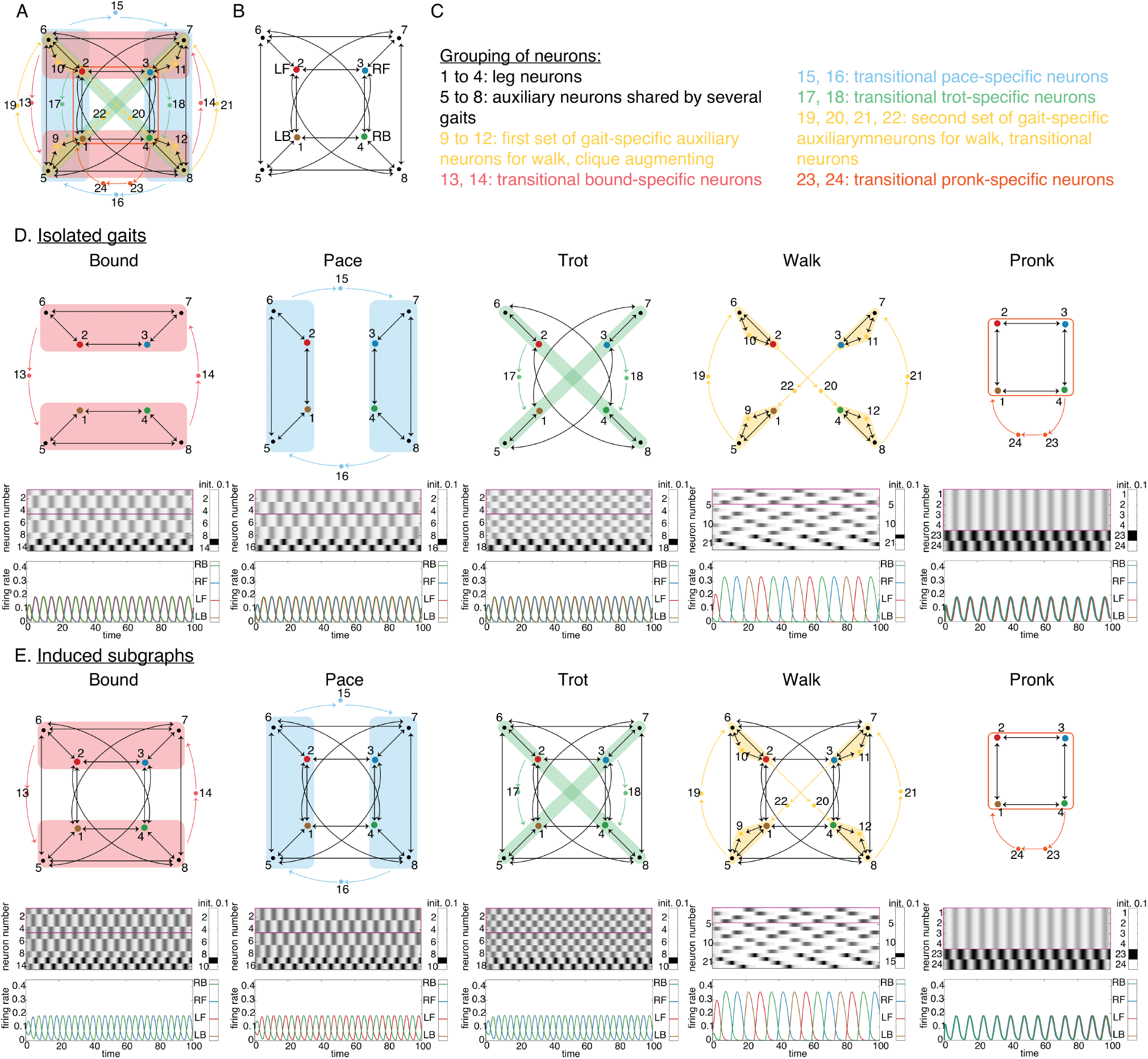
Simulation of five gaits isolated (as opposed to embedded within full network), and as *induced* subgraphs of 5-gait network. Notice that the dynamics of isolated gaits and induced gaits are the same. All simulations shown here were done with *ε* = 0.25, *δ* = 0.5. (A) 5-gait network. (B) Induced subgraph of the 5-gait network nodes 1 - 8. These nodes are shared by two or more gaits. (C) Grouping of neurons by type. Types are assigned on the basis of function, and color coded by gait. (D) Graphs, grayscale and rate curves of all isolated gaits. These are not induced subgraphs. (E) Graphs, grayscale and rate curves of induced subgraphs for all gaits. The attractors corresponding to each gait are still perfectly preserved.

#### Good parameters

A common drawback of many models is that they may require very fine-tuned values of their parameters to perform as expected. To see if this issue arises for the quadruped gait network, we assess the robustness of the network in Figure 4C. Individual gaits were designed as a cyclic union (Thm. 2) of building blocks (Fig. 3) whose FP(*G*) is parameter independent [45], meaning that its FP(*G*) set is preserved across the legal range of parameters 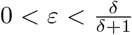. Also, since cyclic unions of parameter independent components are also parameter independent, all isolated gaits are parameter independent as well. Indeed, it is a simple consequence of Theorem 2:

**Corollary 1**. Let *G* be a cyclic union of component subgraphs 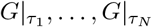. Then FP(*G*) is parameter independent if and only if 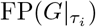 is parameter independent for all *i* ∈ [*N*].

*Proof*. Suppose *σ* ∈ FP(*G, ε, δ, θ*) for all *ε, δ, θ* in the legal range. By Theorem 2, 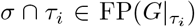 for all *ε, δ, θ* in the legal range and all *i* ∈ [*N*], meaning that 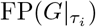 is also parameter independent. The other implication is analogous.

This implies that the FP(*G*) set does not vary with parameter changes within the legal range, and so we can anticipate the gait attractors to be present in each individual gait network for a wide range of parameter values.

Indeed each gait, *isolated*, turns out to be very robust. The same unfortunately cannot be said about the glued five-gait network, since it is not truly a cyclic union, but rather a gluing of cyclic unions, for which we do not have theoretical results yet. This begs the question: how sensitive is the network to the choice of *ε, δ, θ* parameters? Are the five attractors still present, and easily accessible, if we vary the parameter values?

To answer this question, we tried to reproduce the simulations of Figure 4B using different *ε, δ, θ* parameters. First, we observed how the attractor corresponding to each gait behaved under several values of *ε, δ* in the five-gait network. We found that there were three different possibilities: either the attractor is *preserved*, meaning that it is identical in qualitative behavior to the attractors in Figure 4B (up to a change in amplitude or period); *corrupted*, indicating that the auxiliary neurons corresponding uniquely to that gait are still active and firing cyclically, but the leg nodes have lost their symmetry; or *lost*, when the auxiliary neurons corresponding uniquely to that gait are not firing cyclically (this usually translated into a gait becoming a different attractor, which could either be another gait, a stable fixed point or a chaotic-looking attractor).

An example of each of these outcomes is exemplified in Figure S5A, where only the last 200 time units of the simulation are shown, for clarity. In the left example, the network was initialized to the bound gait, and the gait is perfectly preserved. In the middle plot, the network was initialized to the pronk gait, and even though the auxiliary neurons corresponding to pronk are still active, the leg neurons lost their symmetry towards the end. In the right plot, the network was initialized to the trot gait, but near the end of the simulation the network settles into a corrupted form of pronk, as indicated by the auxiliary neurons of pronk being active. The results of all simulations are summarized in Figure S5B. Green denotes preserved attractor, yellow denotes corrupted attractor and red denotes lost attractor. The intersection of the green dotted regions is the range in which all gaits are preserved: 0.05 *< ε <* 0.23, 0.2 *< δ <* 0.6, for at least the first 500 units of time. These are the “good parameters”.

We then did the same analysis but varying the *θ* parameter only. Figure S5C shows examples of what can happen in this case. In the first example, the network was initialized to the walk gait, and the gait is perfectly preserved. In the second example, on the other hand, the network was initialized to the pace gait, but immediately went into pronk. Figure S5D shows the same summary of results, but carried out for constant values *ε* = 0.14, *δ* = 0.40, and varying *θ*. These values were arbitrarily chosen from the 0.05 *< ε <* 0.23, 0.2 *< δ <* 0.6 good parameters from above.

In conclusion, we have computationally found that all the attractors corresponding to the five gaits in our network are preserved for all 0.05 *< ε <* 0.23, 0.2 *< δ <* 0.6 and 0.5 *< θ <* 2.5, for at least 500 units of time. Having a range of parameters available raises the question: what role do the parameters play in modulating the characteristics of the gaits such as period and amplitude? These are important questions because period and amplitude in firing rates affect muscle tension [69, 70].

#### Parameter modulation

Next, we compute the period and amplitude of each gait within the five-gait network, using the good parameters from Figure S5. To do so, we ran a single simulation for *T* = 500 time units and then we used the MATLAB function findpeaks to find the location and value of the peaks of a single rate curve. Examples of this can be seen in Figure S6I. The black bars in the rate curves show the amplitude, and the black triangles mark the time of the peaks, both calculated using only the first leg (*x*_1_(*t*) - LB). We found these peaks only during the last half of the simulation to avoid taking into account the time before settling into the attractor. To find the periods we subtracted all the consecutive peak locations. We had to get rid of the first and last period and amplitude data point because they became corrupted by chopping the curve. Then we averaged the periods and peak values to obtain a single period and a single amplitude per simulation. This is the value shown in Figures S6A-H.

**Supplementary Figure S5:**
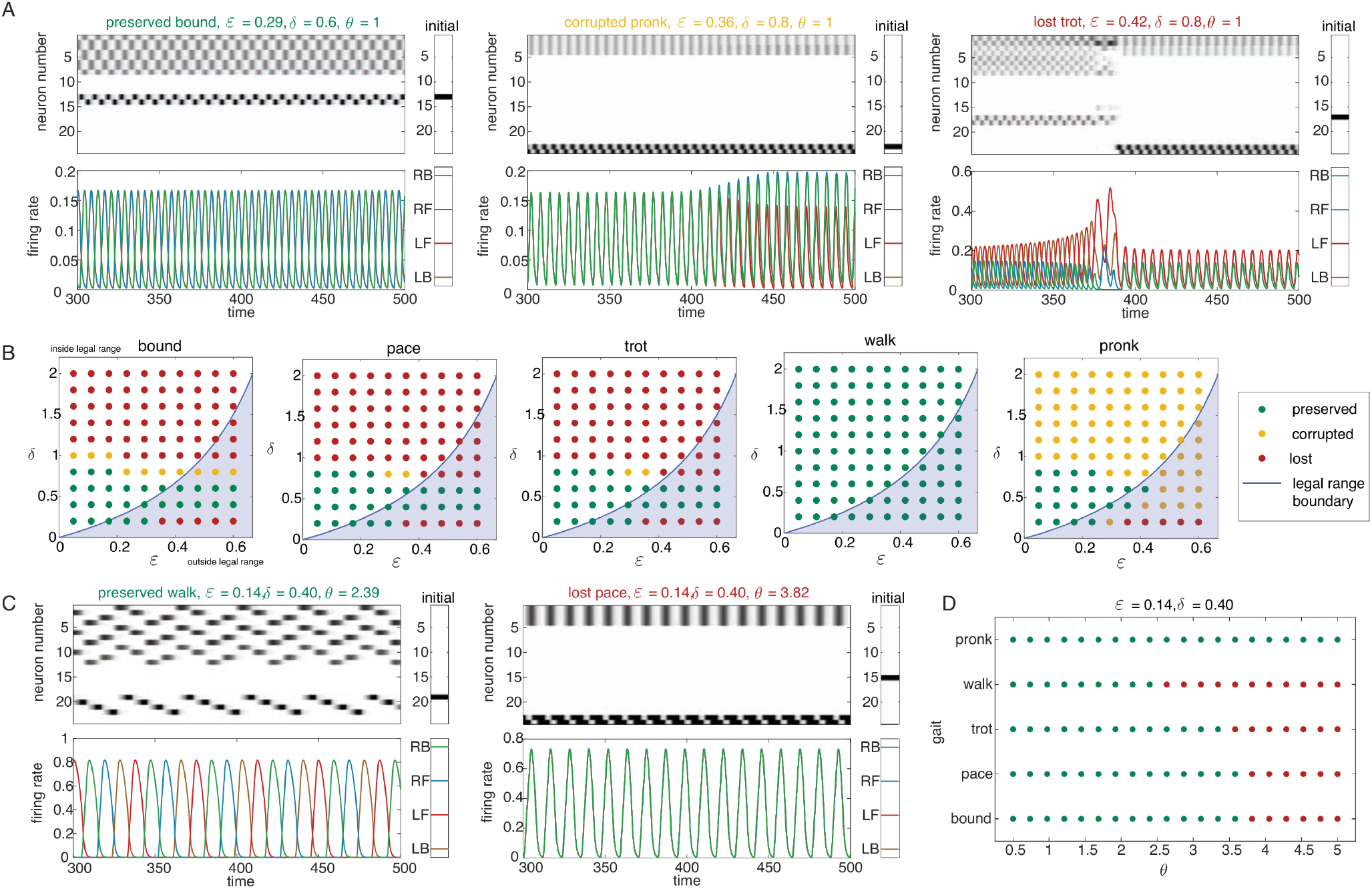
Gaits survival under several values of *ε, δ, θ*. All plots were generated with a running time of 500 time units. (A) Three examples of what can happen to an attractor under three different parameter values. Left: the network was initialized on the bound gait, and the gait is perfectly preserved. Middle: the network was initialized on the pronk gait, and near the end of the simulation, the auxiliary neurons corresponding to pronk are still active, but the limb neurons lost their symmetry. Right: the network was initialized on the trot gait, but near the end of the simulation the network settles into a corrupted form of pronk, as indicated by the auxiliary neurons of pronk being active. (B) Gaits are classified according to whether the attractors are either preserved (green dots), corrupted (yellow dots) or lost to a different attractor (red dots) when *ε, δ* vary. All the dots were generated using *θ* = 1 fixed. (C) Two examples of what can happen to an attractor under two different *θ* parameter values. Left: the network was initialized on the walk gait, and the gait is perfectly preserved. Right: the network was initialized on the pace gait, but immediately went into pronk. (D) Gaits are classified according to whether the attractors are either preserved (green dots), corrupted (yellow dots) or lost to a different attractor (red dots) when *θ* varies. All the dots were generated using *ε* = 0.14 and *δ* = 0.40.

There, each point corresponds to a single simulation. The color of the point corresponds to a given gait, as specified in the label of Figure S6A. Note that, since bound, pace and trot have virtually the same structure, the data point for these are overlapped in Figures S6A-H. We also show cross sections for Figures S6A,D. The blue shaded corner of the 3-dimensional plots corresponds to values outside the legal range. In Figures S6A-F we are varying only *ε* and *δ*, with a fixed value of *θ* = 1. In Figures S6G-H, we use the fixed values of *ε, δ* in the slices above to vary the value of *θ*.

Notice that for fixed values of *ε, δ, θ*, period decreases with *ε* and *δ*; whereas amplitude increases for walk, and decreases for all other gaits. Period is seen to be invariant to *θ* changes, and amplitude increases with increasing values of *θ*. It is known that an increased firing rate produces an increase in muscle tension by summation of more frequent successive motor contractions [69, 70]. This implies that in our model, period would correspond to how fast/slow muscles contract and relax (bigger period means muscles contract and relax faster). The period therefore is controlling how fast the movement is executed, that is, the speed of the gait. Amplitude of firing rate corresponds to tension in muscle, because the more action potentials, the more muscle units are activated.

**Supplementary Figure S6:**
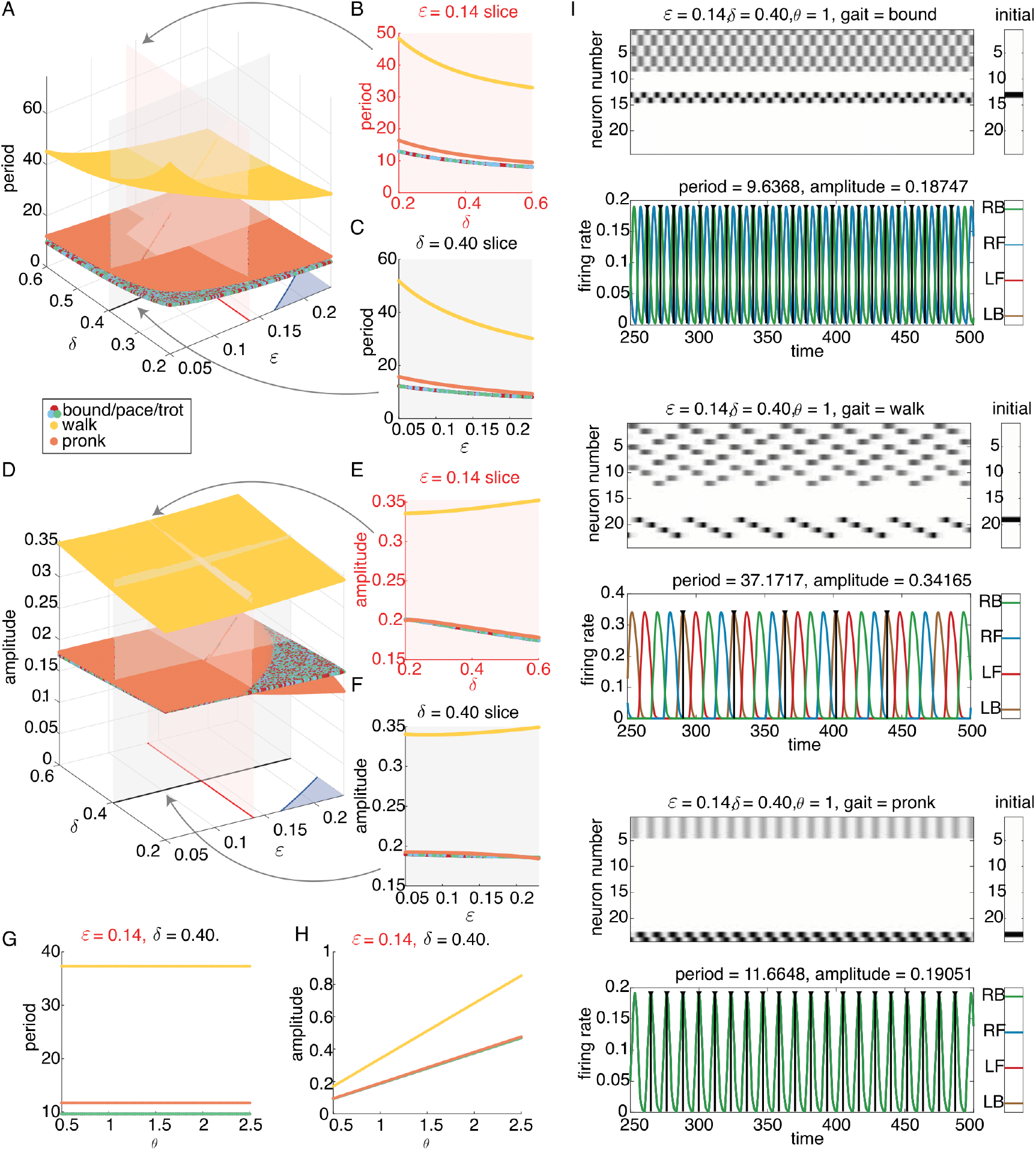
Amplitude and period of the gaits in the 5-gait network (Figure 4C), as controlled by *ε, δ, θ*. Gaits are color coded. Values were found using the last half (once the attractor has settled) of a simulation with running time *T* = 500. (A) Values of period for a (0.05, 0.23) × (0.2, 0.6) grid of (*ε, δ*) values and all gaits. The legal range boundary 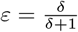 is shown in blue in the (*ε, δ*)-plane. The values above the blue shaded area are outside the legal range, but the gait attractors are preserved nonetheless. The data points for bound, pace and trot are superimposed. (B) Cross section of plot in panel A corresponding to *ε* = 0.14. (C) Cross section of plot in panel A corresponding to *δ* = 0.40. (D) Values of amplitude for a (0.05, 0.23) × (0.2, 0.6) grid of (*ε, δ*) values and all gaits. The legal range boundary 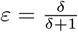 is shown in blue in the (*ε, δ*)-plane. The values above the blue shaded area are outside the legal range, but the gait attractors are preserved nonetheless. The data points for bound, pace and trot are superimposed. (E) Cross section of plot in panel D corresponding to *ε* = 0.14. (F) Cross section of plot in panel D corresponding to *δ* = 0.40. (G) Values of period for the cross sections in panels B and C (*ε* = 0.14, *δ* = 0.40) and varying values of *θ*. (H) Values of amplitude for the cross sections in panels E and F (*ε* = 0.14, *δ* = 0.40) and varying values of *θ*. (I) Three examples of how period and amplitude were computed. The black bars show the amplitude and the black triangles mark the time of the peaks, both calculated using only the first limb (LB).

But while we have showed that there is a fairly wide range of parameters under which our network behaves as expected, and where we can modulate the properties of the firing rates, there is still the assumption that all synaptic connections are perfect and identical: either −1 + *ε* or −1 −*δ*. Is our network robust to noise in the synapses? What about the inputs? Are transitions still possible in the presence of noisy connections?

#### Robustness to noise

To explore these questions, we added uniform noise to *W* and *θ* and computationally tested whether the attractor was lost. A gait is said to be lost when the attractor corresponding to said gait degenerates into a different attractor, which is clear from which auxiliary neurons are active. We did not measure if the attractor was corrupt or not, because any amount of noise will break the symmetry of the legs.

**Supplementary Figure S7:**
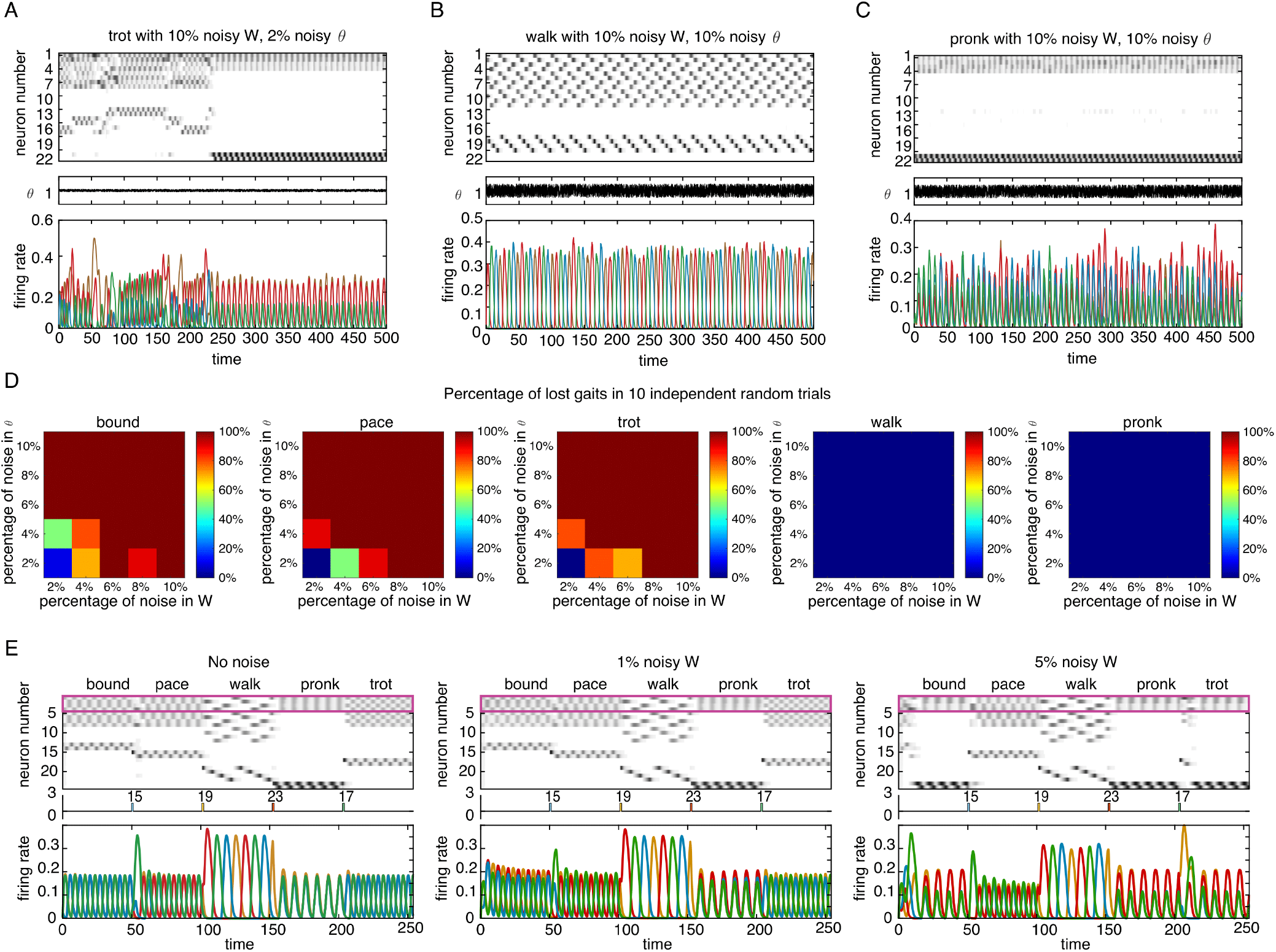
Percentage of lost gaits in noisy 5-gait networks. All simulations used to quantify the failures were done with *ε* = 0.25, *δ* = 0.5 and noise time-scale for *θ dt* = 0.1. We did not measure if the attractor was corrupt or not, because any amount of noise will break the symmetry of the limbs. (A) Example of lost trot. The network chaotically jumps between different attractors, until it settles in pronk. (B) Example of a very noisy network, where walk is not lost, but it is corrupt. (C) Example of a very noisy network, where pronk is not lost, but it is corrupt. (D) Percentage of lost gaits in 10 trials, for several percentages of noise in *W* and *θ*. A trial corresponds to a single matrix, which was used to test all gaits once. (E) Transitions for two values of noise in *W*. Transitions are robust at 1% added noise, but past 5% gaits are commonly lost to pronk and thus transitions are not successful.

This was done in the same way as we did for the TLN counters. We ran a single noisy simulation and observed if the attractor was lost or not. Examples of the behaviors we observed can be found in Figure S7A-C. In A, the network was initialized to trot, but it chaotically jumps between different attractors, until it settles into pronk. In B, the network was initialized to walk, and it remains there, as well as it can. This means, the legs lost their symmetry, but the high-firing neurons are the same as in walk, in the same order. This classifies as not lost in our analysis. In C, we observe the same situation but with pronk. Here, the asymmetry of the legs is more notable. These are wobbly gaits.

Figure S7D summarizes our findings. We simulated each gait 10 times, and then calculated the percentage of lost gaits in those 10 trials, for varying percentages of noise in *W* and *θ*. The color of the pixel in Figure S7D represents the percentage of lost gaits out of these 10 trials. A trial corresponds to a single noisy pair (*W, θ*), which was used to test all gaits once. All simulations used to quantify the failures were done with *ε* = 0.25, *δ* = 0.5 and *θ*-noise with *dt* = 0.1.

In around 90% of trials, gaits were not lost at 2% noisy *W* and *θ*, indicating that the perfect binary synapses of CTLNs are not necessary for this network to produce the desired gaits. Larger levels of noise, however, degrade the network’s performance significantly. Interestingly, at bigger levels of noise, when gaits were lost, the network most commonly settled into the pronk gait. Pronk also happens to be one of the two most robust gaits (see pronk in Figure S7D). This supports our suspicion that pronk’s basin of attraction is bigger. This is perhaps not surprising given how different the construction of pronk is (as well as walk) from the three less robust gaits: bound, pace, and trot. This raises the question of what would happen to the robustness of the network if there were no pronk or walk, so that the network is perfectly symmetric.

Finally, we also tested the ability of the network to transition under 1% and 5% noise in *W*, as shown in Figure S7E. Transitions are robust at 1% added noise, but past 5% gaits are commonly lost to pronk and thus transitions are not successful.

### A.4 Proofs of Section 2.5

**Definition 5**. We say that (*W, b*) is a *competitive* TLN on *n* neurons if *W*_*ij*_ *≤* 0 and *W*_*ii*_ = 0 for all *i, j* ∈ [*n*].

Note that this definition is different from the one in [45], which also requires *b*_*i*_ *≥* 0 for all *i* ∈ [*n*].

**Definition 6**. We say that a TLN (*W, b*) is *non-degenerate* if

- *b*_*i*_ *>* 0 for at least one *i* ∈ [*n*]
- det(*I* − *W*_*σ*_) ≠ 0 for each *σ ⊆* [*n*], and
- for each *σ ⊆* [*n*] such that *b*_*i*_ *>* 0 for all *i* ∈ *σ*, the corresponding Cramer’s determinant is nonzero: det((*I* − *W*_*σ*_)_*i*_; *b*_*σ*_) ≠ 0.

**Lemma 7**. *Let* (*W, b*) *be a competitive, nondegenerate TLN, and suppose b*_*k*_ *≤* 0 *for some k* ∈ [*n*]. *Then x*_*k*_(*t*) → 0 *as t* → *∞. In particular*,

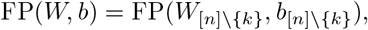

*where* (*W*_[*n*]\ {*k*}_, *b*_[*n*] \ {*k*}_) *is the restricted subnetwork obtained by removing node k from* (*W, b*).

*Proof*. Suppose *b*_*k*_ *≤* 0 for neuron *k*. Then we have that 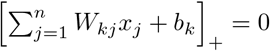, since *W*_*kj*_ *≤* 0 for all *k, j* ∈ [*n*].

Thus 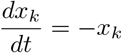, and so *x*_k_ → 0 as *t* → *∞*. Hence at any fixed point, we must have 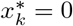. Thus, if *σ* ∈ FP(*W, b*), then *k* ∉ *σ*.

Denote 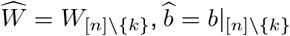. We now show that 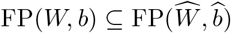, and vice versa. Let *σ* ∈ FP(*W, b*), and let *x*^*\**^ be the associated fixed point. As shown above, 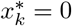. Now let 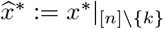. It is easy to check that 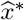 is a fixed point of 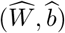. Indeed, for all *i* ≠ *k* we have that

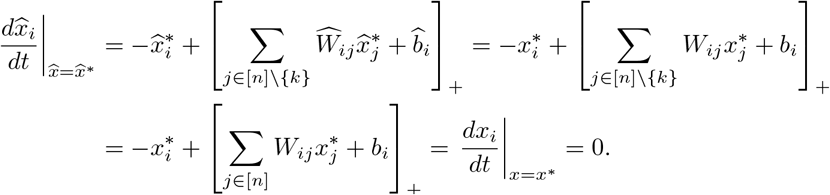

To see that 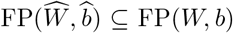, suppose 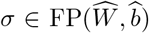 and let 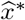 be the associated fixed point. Define *x*^*\**^ such that 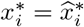 for all *i* ≠ *k*, and 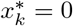. Clearly, supp(*x*^*\**^) = *σ* as well. We now check that *x*^*\**^ is a fixed point of (*W, b*). Indeed, for *i* ≠ *k* we have:

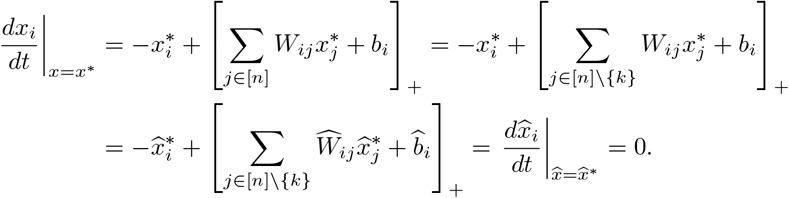

Moreover, since *dx*_*k*_*/dt* = −*x*_*k*_ in (*W, b*), evaluating at *x*^*\**^ we obtain:

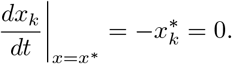

Thus, *x*^*\**^ is a fixed point of (*W, b*), and hence *σ* ∈ FP(*W, b*).

**Theorem 8** (Restatement of Theorem 3). *Consider a TLN* (*W, b*) *where*

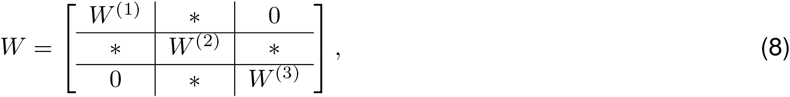

*and*

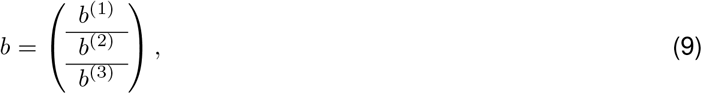

*Proof*. Define the (*i, j*)-th fusion attractor as

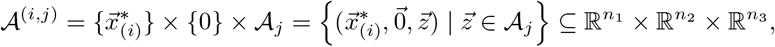

where 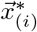 denotes the *i*-th stable fixed point of (*W* ^(1)^, *b*^(1)^).

We will show that 𝒜^(*i,j*)^ is an attractor of (*W, b*), i.e., it is a minimal attracting forward-invariant subset of 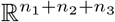.

Let *F*_*t*_(*x*(*t*_0_), *y*(*t*_0_), *z*(*t*_0_)) denote the trajectory at time *t* given an initial condition at *t*_0_, so that

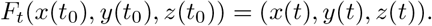

#### Forward invariance

We must prove that for any initial condition (*x*(*t*_0_), *y*(*t*_0_), *z*(*t*_0_)) ∈ 𝒜 ^(*i,j*)^, the trajectory remains in 𝒜^(*i,j*)^ for all *t ≥ t*_0_.

First, note that because *b*^(2)^ *≤* 0, and *W* is competitive, everything inside the nonlinearity is 0 for layer 2 and the dynamics reduce to exponential decay. Since *y*(*t*_0_) = 0 already, it follows that *F*_*t*_(*y*(*t*_0_)) = 0 for all *t ≥ t*_0_.

Next, note that layer 1 only receives inputs from layer 2 and itself. Since *F*_*t*_(*y*(*t*_0_)) = 0, there are no external inputs to layer 1 that can kick the activity out of the initial fixed point of (*W* ^(1)^, *b*^(1)^), and thus 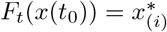 for all *t ≥ t*_0_.

Finally, using the same argument as above (no input from external layers) and noting that *z*(*t*_0_) ∈ 𝒜_*j*_ (which is an attractor of (*W* ^(3)^, *b*^(3)^)), we get that *F*_*t*_(*z*(*t*_0_)) ∈ 𝒜_*j*_ for all *t ≥ t*_0_.

Thus, we have shown that 𝒜^(*i,j*)^ is forward-invariant.

#### Attracting neighborhood

Since 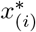 and 𝒜_*j*_ are both attractors, they each have attracting neighborhoods 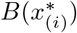 and *B*(𝒜_*j*_). Moreover, since *b*^(2)^ *≤* 0, we have *F*_*t*_(*y*(*t*_0_)) → 0 as *t* → *∞* for all initial conditions, by Lemma 7.

We claim that 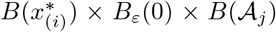 is an attracting neighborhood of 𝒜^(*i,j*)^ for some suitably chosen *ε*. Indeed, it is possible to choose *ε* small enough that the sum of 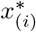 with the *ε* input from layer 2 still falls within the attracting neighborhood 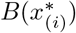. That is, a choice of *ε* is such that the perturbation to the dynamics of layer 1 is small enough for the trajectory to converge to the stable fixed point 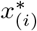. A similar argument applies to the third layer, as the only external inputs it receives come from layer 2, which can be bounded arbitrary close to 0. Taken together, we see that for *t > M, F*_*t*_(*x*(*t*_0_), *y*(*t*_0_), *z*(*t*_0_)) ∈ 𝒜^(*i,j*)^.

#### Minimality

Suppose there exists a smaller subset 𝒜^*′*^ *⊂* 𝒜^(*i,j*)^ that is also an attractor. By the structure of 𝒜^(*i,j*)^, 𝒜^*′*^ must have the form 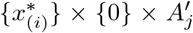 for some proper subset 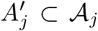. Since the dynamics of layer 3 only involve *W* ^(3)^ inputs in these sets (since layer 2 nodes are 0), the vector field in 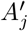 matches that of *A*_*j*_, implying that 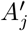 is an attractor of (*W* ^(3)^, *b*^(3)^), smaller than 𝒜_*j*_, contradicting the minimality of 𝒜_*j*_ as an attractor of (*W* ^(3)^, *b*^(3)^).

## Footnotes

1 The upper bound on *ε* is motivated by a theorem in [48]. It ensures that subgraphs consisting of a single directed edge *i* → *j* are not allowed to support stable fixed points, so that activity will flow along the arrows.

2 This is a technical condition first defined in [45] requiring that the relevant determinants of submatrices are nonzero. This condition holds for almost all CTLNs as having a zero determinant is a fine-tuned condition.

